# Structured and flexible representations in medial-frontal cortex support goal-directed navigation

**DOI:** 10.64898/2026.06.09.729603

**Authors:** Peter T. Doohan, Kristopher T. Jensen, Yaqing Chen, Beatriz S. Godinho, Charles D. G. Burns, Chongyu (Xiao) Qin, Josie L. Emery, Ryan J. Cini, Mark E. Walton, Timothy E. J. Behrens, Thomas E. Akam

**Affiliations:** Department of Experimental Psychology, University of Oxford, Oxford, UK; Oxford Centre for Integrative Neuroimaging, University of Oxford, Oxford, UK; Sainsbury Wellcome Centre, University College London, London, UK

## Abstract

Humans and animals plan actions to achieve goals in worlds that are complex and continually changing. While planning is critically dependent on the prefrontal cortex in humans, little is known about its cellular underpinnings. Mechanistic understanding has been limited by a scarcity of controlled animal experiments in which subjects must flexibly plan novel behaviours on every trial. Here we characterise the neural representations and dynamics of mouse medial frontal cortex (mFC) during flexible navigation in structured environments. We trained mice to navigate complex mazes, to goals that changed location on every trial. Optogenetic silencing established that mFC was necessary for efficient navigation. mFC activity was dominated by two factorised components: (i) a structured representation of subjects’ position within the maze that formed an efficient code for behavioural trajectories, and (ii) a flexible representation of the shortest path-distance to the current goal. Both representations oscillated within local field potential (LFP) theta cycles, processing from further to closer to the goal at a systematic offset. These data suggest a computation in which mFC evaluates possible futures by their distance-to-goal to update a structured behavioural policy.

## Introduction

Planning sequences of actions using knowledge of the world’s structure is amongst our most important and complex cognitive functions. Evidence from human neuroscience has implicated the frontal cortex in planning, with lesion [1–4] and fMRI [5, 6] studies emphasising the anterior medial portions as critical. Lesions and inactivations in the rodent homologue, prelimbic medial frontal cortex (mFC), similarly impair goal-directed behaviour and promote habitual responding [7, 8].

Despite the identification of key brain regions involved in planning, little is known about its cellular underpinnings. In mFC, single unit recordings have identified correlates of value [9], task states [10–12], and actions [13] during decision making, but the repetitive nature of most laboratory tasks has made it difficult to concretely link this activity to planning mechanisms. In the hippocampal formation, which shares rich reciprocal connectivity with mFC [14–16], recordings have uncovered internally generated sequential dynamics that represent possible future paths, both during theta sweeps [17, 18] and sharp wave ripples [19]. But whether and how this activity contributes to planning is unclear.

Diverse algorithms have been proposed to underlie biological planning, including sequential search over simulated action sequences [20], representations that embed long-range relationships between states [21, 22], and attractor dynamics that infer desirable futures [23]. Despite this algorithmic diversity, all planning mechanisms share two common components. First, a representation of the world’s structure that constrains future behaviour, and second, a mechanism for flexibly interacting with this representation to determine which actions satisfy current goals. Characterising the representations and dynamics that implement these components will constrain how brains plan. Progress on this front requires new behavioural paradigms [24] that engage flexible, non-repetitive behaviours in structured and dynamic environments — both to recruit planning mechanisms and decorrelate key decision variables.

Here we characterise the neural representations and dynamics of mFC during flexible navigation in structured environments, from which we outline a candidate neural mechanism for making goal-directed decisions.

Using a recently developed behavioural assay [25], we trained mice to navigate to visually-cued goals on complex mazes. Mice learned to navigate flexibly and efficiently to goals that changed on every trial, using knowledge of maze structure over simple heuristics. Optogenetic silencing of mFC reversibly shifted goal-directed strategies towards habitual responding, establishing its causal role in ongoing behaviour. Silicon probe recordings revealed two complementary representations, encoded by largely distinct populations of neurons. The first captured structure that was invariant across trials, comprising cells tuned to conjunctions of current location and movement direction. This representation had consistent low-dimensional structure that reflected both structural features of the maze and the statistics of behaviour, forming an efficient code for behavioural trajectories. The second representation was of a decision variable: it comprised a population code for shortest-path distance-to-goal that flexibly updated as goals changed location. These two codes interacted dynamically on both a behavioural timescale, and within individual theta-cycles, where the position and distance codes swept from further to closer to the goal at a systematic temporal offset. Together these findings suggest a decision-making algorithm in which theta-sweeps in the hippocampal formation read out possible futures into mFC, that are evaluated by their distance-to-goal, to shape a structured and efficient representation of a behavioural policy.

## Results

### Goal-directed navigation in complex mazes

We trained mice to navigate an elevated grid-maze comprising 49 towers arranged in a 7-by-7 grid, housed in a dark enclosure (Figure S1a). Each tower contained a stimulus LED, a speaker, and a reward port. Adjacent towers were connected by removable walkways to create complex maze layouts. On each trial, a randomly chosen tower (the ‘goal’) was cued by illuminating its LED and playing a brief white noise (500 ms). Subjects navigated to the cued goal and received a water reward, followed by an intertrial interval (ITI; 4-8 s; Figure 1a-b).

**Figure 1:**
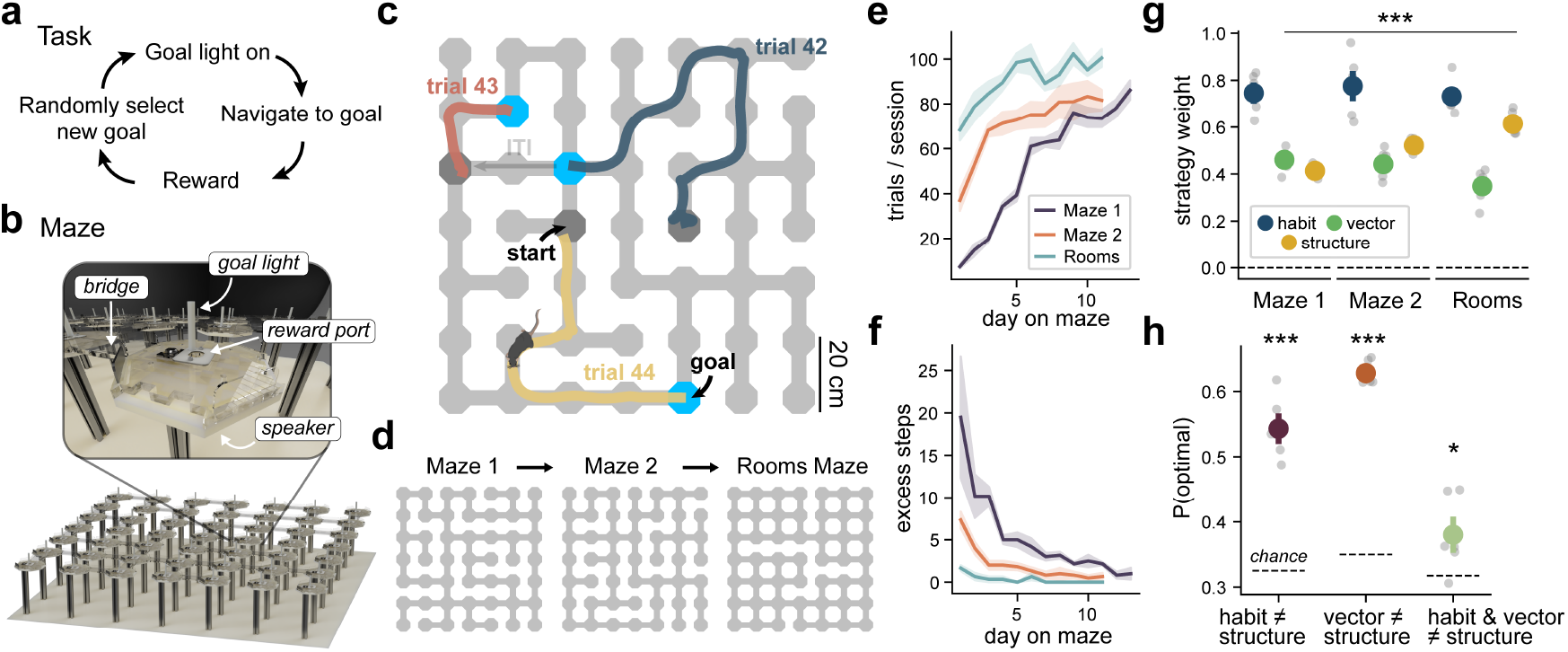
Mice learn to flexibly navigate complex mazes. **(a)** Task design: mice performed a goal-directed navigation task embedded in complex mazes. On each trial, a goal location was randomly selected and cued with a brief auditory and sustained visual cue. After successfully navigating to the goal subjects received a water reward, followed by a brief inter-trial-interval (ITI) before the next goal was cued. **(b)** Maze apparatus: mice navigated a programmatically controlled grid maze apparatus consisting of 49 elevated towers connected by removable walkways (bridges), fully enclosed in a dark enclosure (not shown). **(c)** Navigational trajectories from three consecutive trials. Note: subjects can freely move after reward consumption during the ITI period, so the start locations (dark grey locations) can differ from the previous trial’s goal (blue locations). **(d)** Subjects were exposed to a series of 3 maze configurations (for 11-13 days each). Mazes 1 & 2 were optimised to decorrelate shortest-path and Euclidean distances between towers. The Rooms Maze was designed as four fully connected “rooms” connected by single walkways. **(e)** Trials completed per session increased with experience on each maze and across mazes. **(f)** Subjects deviated less from the optimal paths between start and goal locations (took fewer excess steps) with experience on each maze and across mazes. **(g)** Mixture-of-strategies model fit to subject’s navigational decisions. All strategies: habitual choice (“habit”; repeat most likely action given current and prev. location, independent of current goal), vector navigation (“vector”; reduce Euclidean distance to goal) and structure-based (optimal) navigation (“structure”; reduce shortest path distance to goal) significantly predicted behaviour. **(h)** Probability subjects chose optimal (structure-based) actions at decision points (towers with *>* 2 connected bridges), restricted to decisions where: i) habit alone disagreed with structure (purple); ii) vector alone disagreed with structure (dark orange); iii) or both disagreed with structure (light green). Dashed lines mark chance. All panels: error bars and shaded errors represent SEM across subjects; individual subject values plotted as grey markers. *t*-test *>* 0 or chance: *p<.05, ***p<.001.

Over a session, mice continually navigated between different start and goal locations. Each trial constituted a new navigation problem, solved by generating a novel navigational trajectory (Figure 1c). After learning the cue-reward association during pretraining (Figure S1d-e), mice performed this task on a series of 3 different mazes (Figure 1d; Figure S1f). Mazes 1 & 2 were optimised to maximally decorrelate the Euclidean and shortest-path distances between maze locations (Figure S1c). Maze 3 was designed as four fully connected “rooms” adjoined by single links (Rooms Maze), motivated by work on hierarchical reinforcement learning [26]. The number of trials completed per session increased with experience, both within each maze and across mazes (Figure 1e; linear mixed model: day: *β* = 8.4, *z* = 16.6, *p <* .001, maze×day: *β* = −2.1, *z* = −8.2, *p <* .001). Congruently, paths taken to the goals became more efficient over days and across mazes (Figure 1f; linear mixed model: day: *β* = −1.6, *z* = −8.66, *p <* .001, maze×day: *β* = 0.5, *z* = 5.6, *p <* .001).

To quantify how different strategies contributed to behaviour, we modelled choices during navigation with a mixture-of-strategies model that combined: (i) vector navigation (reduce Euclidean distance to goal); (ii) habitual action selection (repeat the most likely action given current and previous location); and (iii) an optimal structure-based strategy (reduce the shortest-path distance to goal; Figure S1g). The fitted weights revealed that all strategies contributed significantly to behaviour across subjects (Figure 1g, Figure S1h; one-sample *t*-tests *>* 0: *T* (5) *>* 12, *p <* .001, all strategies, all mazes).

To summarise their relative contributions, we measured the rate of optimal choices at decision points when habit, vector, or both strategies prescribed suboptimal actions (Figure 1h). When only one alternative strategy was suboptimal, subjects strongly favoured the optimal action (paired *t*-tests vs. chance; habit ≠ structure: *T* (5) = 11.6, *p <* .001; vector ≠ structure: *T* (5) = 33.8, *p <* .001), establishing structure-based navigation as a dominant driver of choice. However, when both habit and vector strategies were suboptimal, optimal choice rates only just exceeded chance (*T* (5) = 2.8, *p* = .037) — highlighting the residual influence of lower-level strategies in certain conditions. Thus, mice learned to navigate complex mazes flexibly and efficiently by drawing on knowledge of maze structure.

### Silencing mFC impairs goal-directed navigation

Human medial prefrontal lesions strongly impair goal-directed behaviour [1, 3, 4]. However, most rodent laboratory tasks used to study goal-directed behaviour do not require mFC after learning [27– 29, 8, 30]. We therefore asked if mFC was necessary to flexibly navigate structured mazes.

To probe the contribution of mFC to ongoing behaviour, we silenced mFC optogenetically during navigation. We trained mice expressing either the inhibitory opsin stGtACR2 [31] (opto group, *n* = 7) or a control construct (control group, *n* = 8) in mFC (Figure 2a&b, Figure S3a). Once task performance on a maze plateaued (8 days for Maze 1 & 5 days for Maze 2; Figure S2a), blue light (473 nm) was delivered bilaterally through an optic fibre to mFC throughout the navigation period on a randomly selected 25% of trials (light-ON trials). Control experiments verified that this approach effectively silenced mFC neurons (Figure S2). After 11–12 days of stable (expert) behaviour, subjects were exposed to a new maze and the protocol was repeated.

**Figure 2:**
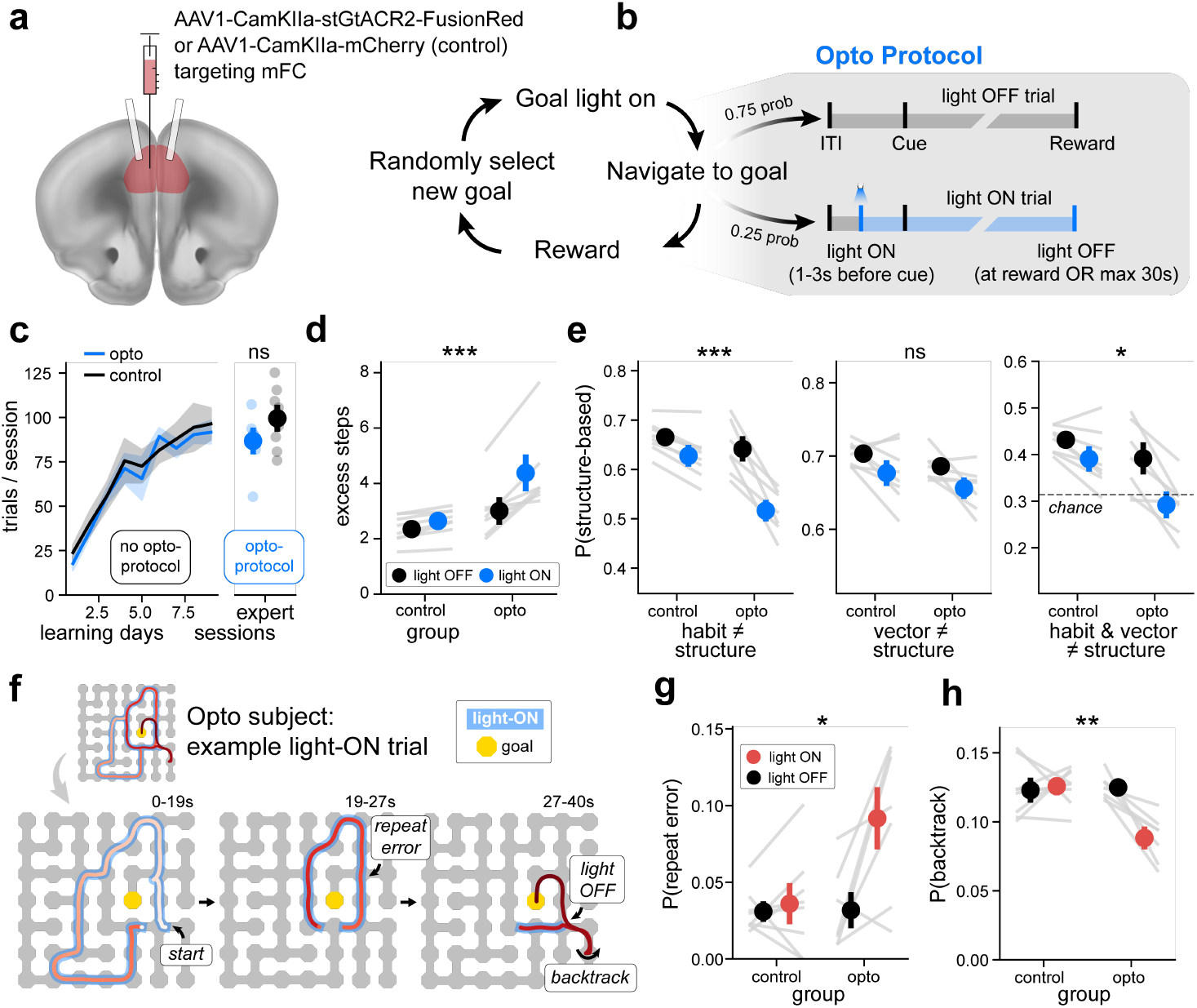
Silencing mFC shifts goal-directed navigation towards habitual action selection. **(a)** Schematic illustration of optogenetic approach using brain slice from Allen CCFv3 MRI reference atlas. The inhibitory opsin “stGtACR2” was used to silence mFC activity during navigation. **(b)** Task structure adapted for optogenetic manipulations. During expert behaviour, blue light (473 nm) was delivered to mFC via a tethered optic fibre on random 25% of trials to silence neural activity in opsin-expressing subjects. **(c)** Opto and control groups performed a similar number of trials over learning (no manipulation) and expert behaviour (w/ opto protocol). Data from the 1^*st*^ Maze. **(d)** Silencing mFC made paths to goal less efficient. Plotted: average number of excess steps per trial in opto and control groups across light-ON (blue) and light-OFF (black) conditions. Individual subject values plotted as connected grey lines between conditions. **(e)** Silencing mFC significantly increased the likelihood of making navigational errors when the optimal choice (structure-based) conflicted with either habitual action selection (left) or both habitual action selection and vector-navigation (right), but not errors when optimal choices conflicted with vector-navigation alone (middle: structure ≠ vector, 3-way ANOVA: *F* (1, 39) = 1.28, *p* = .132). Plots formatted same as panel d. **(f)** Example light-ON trial trajectory in an opsin expressing subject, broken down into three consecutive segments (left to right). Behavioural trajectory coloured by time in trial (white: start, dark red: end), blue highlights indicate light-ON period (max 30 s), goal location coloured yellow. **(g)** Silencing mFC made subjects more likely to commit the same error multiple times. Errors defined as choices that increased distance to goal after it was previously decreasing. See example “repeat error” in panel f (middle). Plotted: mean repeat error probability across conditions, formatted as in panel d. **(h)** Silencing mFC made subjects less likely to backtrack after an error. See an example “backtrack” in panel f (right). Plotted: mean probability of backtracking 1-2 steps after a navigational error across conditions, formatted as in panel d. All panels: Error bars and shaded error represent SEM. Two or three way ANOVA interaction term: *p<.05, **p<.01, ***p<.001.

Both opto and control groups performed similar numbers of trials-per-session during learning (Figure 2c, Figure S3a; linear mixed model group×day: *β* = 0.15, *z* = .14, *p* = .887), and after learning (Fig- ure 2c, Figure S3a; *t*-test: *T* (13) = 1.48, *p* = .162). Additionally, both groups started moving towards the goal shortly after the goal cue (Figure S3g), suggesting that mFC inhibition did not decrease general motivation or on-task behaviour. Strikingly however, the opto group took significantly less efficient paths to goal under mFC-inhibition (Figure 2d; mixed ANOVA group×light-ON: *F* (1, 13) = 15.91, *p* = .0015, 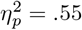). This increase in excess steps reflected an increased probability of making navigation errors when optimal choices conflicted with those prescribed by habitual action selection (Figure 2e; mixed ANOVAs group×light-ON×strategies-disagree. structure ≠ habit: *F* (1, 39) = 15.2, *p <* .001) or both habitual and vector-based strategies (structure ≠ habit or vector: *F* (1, 39) = 3.51, *p* = .034). mFC inhibition was most disruptive when navigating to goals that were hard to reach by vector navigation, or to goals that were located in peripheral, and therefore less frequently visited, parts of the maze (Figure S3d-f).

Impaired goal-directed behaviour during mFC inhibition was exemplified by striking light-ON trials where opto subjects continued to follow habitual routes with apparent disregard for the current goal — sometimes making the same navigational error multiple times (“repeat errors”; Figure 2f, Figure S3c). Indeed, opsin-expressing subjects made significantly more “repeat errors” on light-ON trials (Figure 2g; mixed ANOVA group×light-ON: *F* (1, 13) = 12.7, *p* = .003, 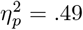) and were less likely to correct errors by turning back (Figure 2h; mixed ANOVA group×light-ON: *F* (1, 13) = 23.1, *p <* .001, 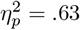). To further quantify how mFC inhibition affected maze navigation, we fitted our mixture-of-strategies model to predict decisions in light-ON and light-OFF trials separately. Comparisons of strategy weights across conditions confirmed that silencing mFC increased habitual navigation (Figure S3h), and significantly decreased structure-based navigation on Maze 2 (Figure S3h). Together these results demonstrate that mFC is necessary for efficient goal-directed navigation. Without mFC, goal-directed behaviour gives way to habitual strategies.

### mFC recordings during flexible navigation

Having established that flexible navigation on the maze requires mFC, we set out to understand the neural representations and dynamics underlying this behaviour. We used silicon probes to record mFC neurons — predominately in the prelimbic and anterior cingulate regions — while mice performed the task (Figure 3a-b; Figure S4a-b). Population firing rates showed a transient peak following the goal cue, and suppression during reward consumption (Figure 3c-d). Individual mFC neurons tiled the different task phases, with some neurons active only during reward consumption or the inter-trial interval (ITI). Many neurons were active during navigation (Figure 3e-f; Figure S4e), which was also marked by an increase in theta (7-11 Hz) and 4-5 Hz LFP power (Figure 3g-h; Figure S4c-d). Next we focus on neural activity during navigation, asking how mFC combines (i) a representation of maze structure and animals’ current behaviour (Figure 4) with (ii) information about the current goal (Figure 5), to select goal-directed actions (Figure 7).

**Figure 3:**
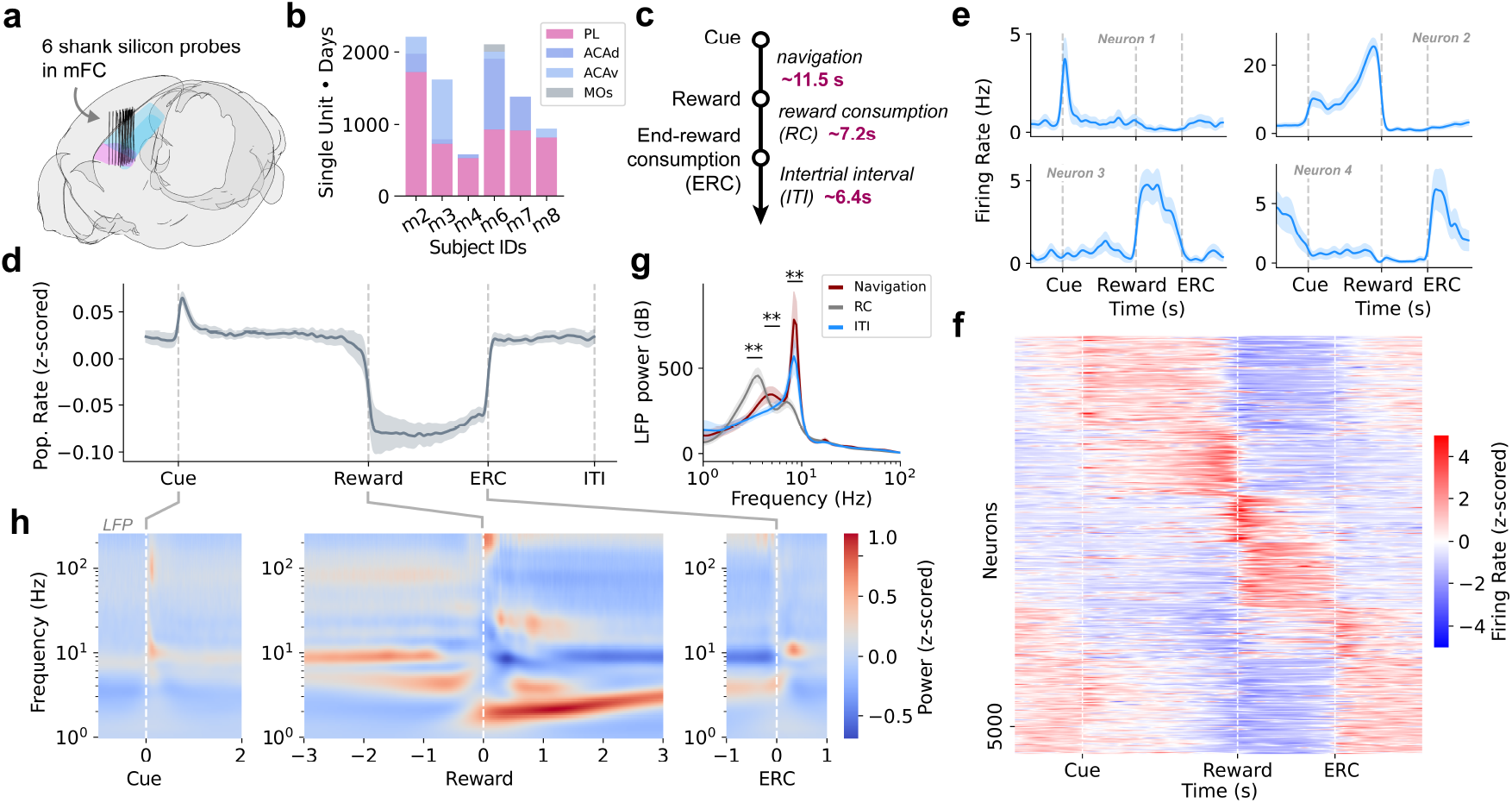
mFC represents task structure. **(a)** Silicon probe tracks reconstructed in Allen CCFv3 space. Probe contacts targeted the left mFC (primarily the prelimbic (pink) and anterior cingulate (cyan) cortical regions). **(b)** Single unit counts in each mFC region across subjects. PL: prelimbic cortex, ACAd: dorsal anterior cingulate cortex, ACAv: ventral anterior cingulate cortex, MOs: secondary motor cortex. **(c)** Trial events (left) and trial phases (right). The median trial phase duration across late sessions is printed next to each trial phase in purple. **(d)** Mean mFC population activity across trial events. **(e)** Example single units tuned to trial events (e.g., cue: top left), and trial phases: navigation (top right), reward consumption (bottom left) and ITI (bottom right). **(f)** Population trial-aligned tuning heatmap demonstrates subpopulations of mFC tuned to the trial phase epochs (navigation, reward-consumption and ITI). Neurons were ordered into 6 agglomerative clusters (using a cross-validated grouping procedure) and ordered by agglomerative cluster group. **(g)** LFP power spectral density in each trial phase epoch demonstrates prominent slow oscillations (2-4 Hz) during reward consumption, strong theta (7-11 Hz) power during navigation and the intertrial interval period and an additional peak at 4-5 Hz during navigation that was not present during the ITI epoch. *t*-tests across trial-phases for each freq. band: **p<.01. **(h)** Mean power spectral density of LFP signal (z-score normalised across frequencies) aligned to cue (left), reward (middle) and end-reward-consumption (ERC, right), highlight broad signal deflections at cue and ERC as well as sustained increases in LFP theta and 4-5 Hz power during navigation before reward. All panels: Shaded error represents SEM.

**Figure 4:**
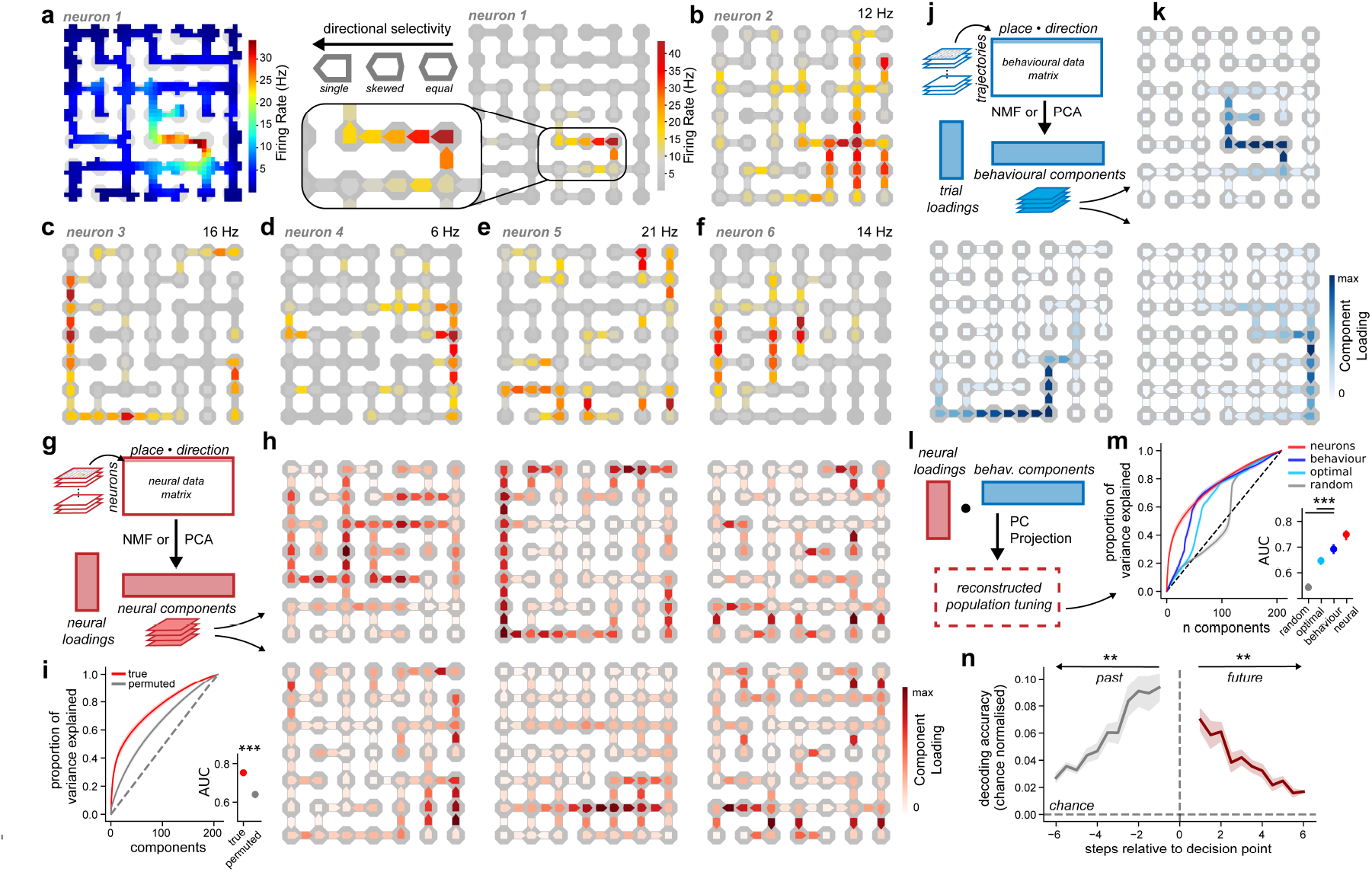
Structured spatial tuning forms an efficient code for behavioural trajectories. **(a)** Neurons selectively fired when moving through maze locations in particular directions. We visualised this with “place-direction” heatmaps that jointly visualise average firing rate (colour) and directional preference (marker shape). Plotted: standard heatmap (left) and place-direction heatmap (right) of the same cell. **(b-f)** Single neurons that fired directionally towards decision points (b), fired directionally along peripheral routes (c-d), from high-centrality locations towards dead-ends (e), and along maze segments with common local structure (f). **(g)** Dimensionality reduction (NMF or PCA) on population place-direction tuning, yields neural components that can be visualised as “place-direction” heatmaps. The “neural data matrix” combines place-direction tuning curves across neurons recorded on a given maze. **(h)** NMF components tended to flow directionally through bottlenecks, around extended routes on the border of different mazes, and between high centrality locations and dead-ends (select components from 10 component NMF performed separately on each maze; see Figure S6a for all components). **(i)** Neural place-direction tuning was lower-dimensional than expected by autocorrelation in behaviour and neural activity. Plotted: cumulative cross-validated variance explained by principal components from PCA on true or permuted population place-direction tuning (left) with area under the curve (AUC) summaries (right). **(j)** Dimensionality reduction (NMF or PCA) on trial-by-trial behavioural trajectories. The “behavioural data matrix” combines trajectories (sequences of occupied place-directions) across trials. **(k)** NMF yielded components tuned to extended action sequences that appeared systematically related to single-neuron tuning and neural NMF components (select component from 10 component NMFs, see Figure S6c for all components). **(l)** To quantify the similarity between low dimensional structure in neural place-direction tuning and subjects’ behaviour we used the behavioural principal components (from PCA) as a basis set to explain variance in neural place-direction tuning. **(m)** PCs derived from subject’s behaviour (blue) explained variance almost as quickly as the neural PCs themselves (red) and were better at explaining neural variance than PCs derived from optimal behaviour (cyan) or random diffusion (grey). Right: Area under curve (AUC) of these cumulative distributions. **(n)** Past and future place-direction occupancy could be decoded from neural activity at decision points, above that implied by subject’s current place-direction (chance), using multinomial logistic regression. *t*-test vs. chance, BH-corrected: **p<.01. All panels: shaded error and error bars represent SEM across subjects.

**Figure 5:**
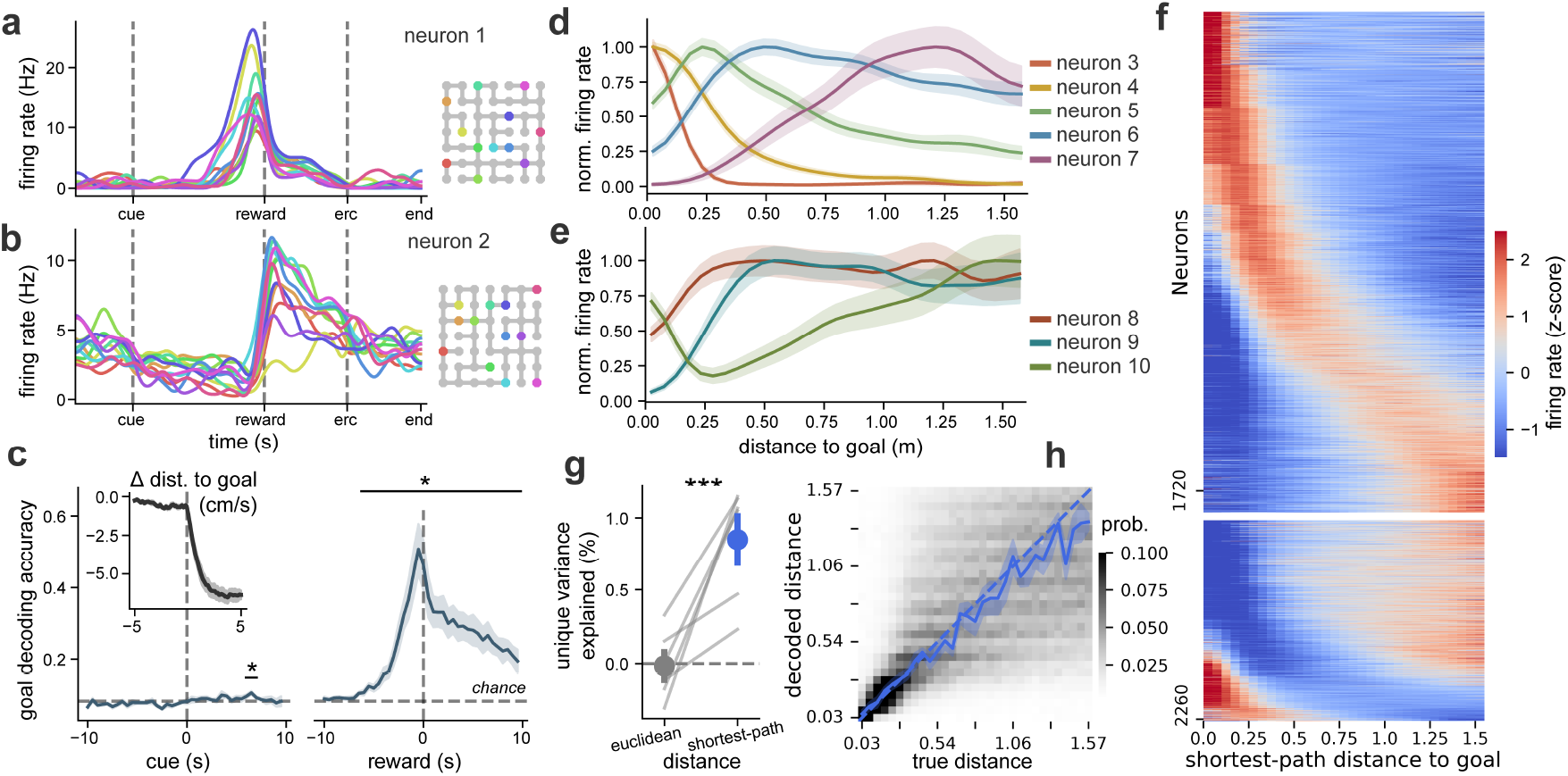
A population code for shortest-path distance to goal. **(a-b)** Single-unit trial-aligned firing rates differentiated between goal locations just before reward (a) and during reward consumption (b). Plotted: trial-aligned activity averaged across trials for the same navigational goal, coloured according to maze legend (right). **(c)** Above-chance goal-decoding at reward time but not at the start of navigation. Allocentric goal decoding accuracy at different time-points relative to trial cue and reward (average decoding across all 12 goal sessions). Left inset: average rate of change of shortest-path distance to goal (Δ Dist. to goal) aligned to cue, indicates that animals were headed to the goal, while goal decoding performance was at chance levels. **(d-e)** Example neurons tuned to particular distances to goal, either firing maximally (d) or minimally (e) at some preferred distance. Neurons normalised by their maximum firing rate, ordered by their preferred distance, and coloured separately. **(f)** mFC population tuning to distance-to-goal. Distance-to-goal tuned neurons were split into “positive” (e.g., neurons in d) and “negative” (e.g., neurons in e) subpopulations (top block and bottom block, respectively), and ordered by their maximum/minimum firing rate from a fitted gamma distribution. **(g)** Shortest-path distance rather than Euclidean distances better explains variance in population distance-to-goal coding. Plotted: mean coefficient of partial deviance (unique variance explained %) for each distance metric. Grey lines represent single subject values. **(h)** Performance of a logistic regression decoder predicting distance-to-goal from neural activity. Background heatmap displays the average decoded distance probability distribution, at each (true) distance-to-goal. The most likely decoded distance (full blue line) closely matched the true distance (dashed-blue line) across distances. All panels: error bars and shaded error represents SEM across trials for single neurons else SEM across subjects. Paired or one-sample *t*-test: *p<.05, ***p<.001.

### Structured tuning to place and movement direction

Frontal cortex is thought to encode spatial information during navigation [13, 32], but the representational principles and computational function are poorly understood. Next we asked: (i) if mFC represented subjects’ position in the maze, and (ii) whether this representation was systematically related to maze structure.

Many mFC neurons had firing fields localised to particular portions of the maze (Figure 4a–left), similar to place cells in the hippocampus [33]. However, the firing fields of mFC cells were often highly direction sensitive: they fired only when animals moved through a particular location in a particular direction (Figure 4a–right). To jointly visualise tuning to place and direction we created “place-direction heatmaps” which indicate average firing rate at a location by colour, and directional selectivity by marker shape (Figure 4a). Consistent patterns of place-direction tuning were observed across subjects and appeared systematically related to the maze structure. Examples included neurons with place-direction tuning to (i) directional movement towards decision points (Figure 4b); (ii) directional movement along the periphery of the maze (Figure 4c-d); (iii) movement from high-centrality locations towards dead-ends; and (iv) movement along maze segments with common local structure (Figure 4f).

To characterise place-direction tuning at the population level, we performed dimensionality reduction on a “neural data matrix” obtained by stacking place-direction tuning curves across neurons recorded on a given maze (Figure 4g). We applied non-negative matrix factorisation (NMF) to extract shared structure across neurons, yielding components that corresponded to directional movement: (i) around central parts of the maze; (ii) through bottleneck locations; (iii) along prominent peripheral routes; and (iv) between high-centrality locations and dead ends (Figure 4h, Figure S6a). We verified that this low-dimensional structure was consistent across subjects (Figure S5b), and was not an artefact of autocorrelation in neurons and behaviour (Figure 4i, Figure S5a; permutation tests: p<.001 for all mazes). Finally, we used representational similarity analysis (RSA) to confirm that significant neural variance could be explained by structural maze features including decision-points, dead-ends, borders, and corners (Figure S5g).

To investigate whether single neurons consistently represented the same abstract structural features, we tracked neurons across mazes with UnitMatch [34]. Across neurons, remapping between mazes was highly non-random (Figure S5f; true vs. permuted match tuning curve correlation; paired *t*-test: *T* (5) = 4.64, *p* = .006). Remapping was not limited to cells responding at the same allocentric location, instead we found neurons consistently tuned to the same structural features across environments. For example, some neurons were consistently tuned to peripheral routes, despite those routes following different borders on different mazes. Others reliably coded for movement between central parts of the maze and dead-ends, despite the locations of those dead-ends changing drastically from one maze to the next (Figure S5e, Figure S6e).

### An efficient code for behavioural trajectories

This structured spatial tuning might represent either where the animals are in the environment, or what they are doing — their current *state* or *policy*. To probe this distinction, we asked whether the mFC place-direction representation was systematically related to the animals’ behaviour, testing i) whether tuning curves reflected the statistics of behaviour across trials, and ii) whether activity predicted future navigational choices on individual trials.

Navigation induces strong correlations among which place-directions co-occur on the same trajectory: passing through one location strongly predicts passing through neighbouring locations in the same direction. Just as sensory regions are thought to efficiently encode the statistics of their inputs [35], an efficient neural code for behavioural trajectories should exploit this low-dimensional structure to reduce redundancy. To characterise the low-dimensional structure of behaviour, we constructed a ‘behavioural data matrix’ whose rows were binary vectors indicating the place-directions visited on each trial (Figure 4j). Using principal component analysis (PCA) we verified that a small number of components explained a large fraction of the variance in the behavioural data matrix (Figure S5c). We visualised this low-dimensional structure using NMF, yielding components that corresponded to trajectory segments on the maze (Figure 4k; Figure S6c).

To test whether neural place-direction tuning reflected the low-dimensional structure of behaviour, we performed PCA on the neural data matrix and the behavioural data matrix separately (Figure S6b&d). We then quantified how quickly variance in the neural data matrix was explained by both neural and behavioural principal components (PCs; Figure 4l). In other words, we asked whether components that explain large amounts of variance in behavioural trajectories across trials also explain a large amount of variance in place-direction tuning across neurons. Strikingly, PCs derived from subjects’ behaviour explained variance in the neural data almost as quickly as PCs derived from neural activity itself (Figure 4m). PCs derived from the empirical behaviour provided a more efficient basis for neural tuning than PCs derived from either a simulated random walk or fully optimal behaviour (Figure 4m, Figure S5d; permutation tests: true behaviour vs. optimal: p<.001, true behaviour vs. random: p<.001, for all mazes). Together, these results show that mFC place-direction tuning constitutes an efficient code for behavioural trajectories.

The preceding analysis demonstrates that mFC represents the average statistical structure of behaviour. Next we asked if mFC contained information about past and future behaviour on a trial-by-trial basis. We used multinomial logistic regression to predict past and future place-directions from neural activity at decision-points, while controlling for information provided by the current place-direction. Neural activity significantly improved decoding accuracy up to 5 steps into the past and future (> 1m; Figure 4n, Figure S5h; paired *t*-tests, BH-corrected *p* < .01, all time-points).

Taken together, these results demonstrate that spatial representations in mFC combine information about (i) the structure of the environment, (ii) the statistics of behaviour, and (iii) the specific past and future trajectory on each trial. Thus, mFC contains information about subjects’ current behaviour or *policy*, more than simply where they are on the maze.

### mFC represents shortest-path distance to goals but not their location

Flexible behaviour requires mechanisms that shape the current policy towards current goals. We therefore asked whether mFC contained a flexible representation that adapted as goals changed.

One candidate would be a representation of the goal itself, as the frontal cortex is widely believed to represent behavioural goals [32, 36, 37, 30, 38]. Indeed, many mFC neurons differentiated between navigational goals around the time of reward and reward consumption (Figure 5a-b), and we could accurately decode navigational goals from mFC activity during these periods (Figure 5c; *t*-tests *>* chance, BH-corrected: *p <* .05). However, goal decoding was at chance levels early in navigation, at time-points when subjects were already *en-route* to the goal — as indicated by the rate of change of distance-to-goal (Figure 5c-inset). In fact, goal decoding in our data matched what would be expected from the place-direction tuning (Figure S7g) rather than an explicit representation of the goal itself. Decoding was not meaningfully improved by the use of data-aggregation techniques (Figure S7h) or more flexible decoders (Figure S7i).

Although goal locations were not explicitly represented, goals did profoundly shape mFC activity: through a population code for shortest-path distance-to-goal. Different neurons fired maximally at different distances-to-goal (Figure 5d-e), such that a population of cells tiled distance-to-goal space (Figure 5f; Figure S8b). Neural tuning was precise at short distances and broader at long distances, with population tuning curves being well fit by a family of gamma-distributions (Figure S8c). Strikingly, this distance tuning generalised across mazes, where individual neurons showed similar tuning profiles across environments with radically different distances between the same pairs of locations (Figure S8e-f).

To test whether Euclidean or shortest-path distance better explained neuronal tuning, we modelled mFC activity using Poisson regression with Euclidean and shortest-path distance basis functions as predictors. We found that only shortest-path distance explained unique variance (Figure 5g; paired *t*-test Euclidean vs. shortest-path: *T* (5) = 4.6, *p* = .006; *t*-test Euclidean *T* (5) = − 0.16, *p* = .88). When repeating this analysis with different distance- or progress-to-goal metrics, variance was best explained by either shortest-path distance or “future path distance” along the actual path taken by the animal — which were highly correlated (Figure S8a). Congruent with tuning patterns across neurons, we could decode shortest-path distance-to-goal from mFC activity using multinomial logistic regression (Fig- ure 5h). Decoding was most precise at shortest distances (< 0.6 m, ≈3 towers), but remained accurate over a wide range of distances (cross-subject average decoding Figure 5h-blue line), even as the decoded probability mass became more broadly distributed (i.e., less precise; Figure 5-heatmap).

Thus, mFC implements a population code for shortest-path distance-to-goal. This provides a putative decision variable: at each decision-point, selecting the upcoming option with the shortest distance-to-goal is sufficient to produce optimal trajectories.

### Factorised representations of place-direction and distance-to-goal

We have shown that mFC represents both the animal’s current place-direction and the associated distance to the trial-specific goal. Yet analysing these variables separately cannot answer important questions about how they jointly determine activity: Do they explain unique variance? Do they factorise or interact non-linearly? Are they represented by the same or different neurons?

Generalised linear models (GLMs) are the standard tool for partitioning neural variance across behavioural features [39]. However, standard GLMs are ill-suited to our task, as the space of behavioural variables and their possible interactions is prohibitively high-dimensional relative to the data collected per neuron [40–43]. To overcome this challenge, we developed a new Neural Embedding GLM (neGLM; Figure 6a-b) that learns a compression of the feature space by combining data across sessions. neGLM comprises two stages: (i) learning a nonlinear neural network embedding that compresses high-dimensional behavioural features into a low-dimensional latent space (Figure 6a), and (ii) a cross-validated Poisson GLM that predicts neural activity in held-out sessions as linear combinations of the learned latent features (Figure 6b). Critically, neGLM shifts the burden of fitting arbitrary high-dimensional tuning curves from a per-neuron GLM to a joint embedding that can be learned over an entire dataset of non-simultaneously recorded neurons.

**Figure 6:**
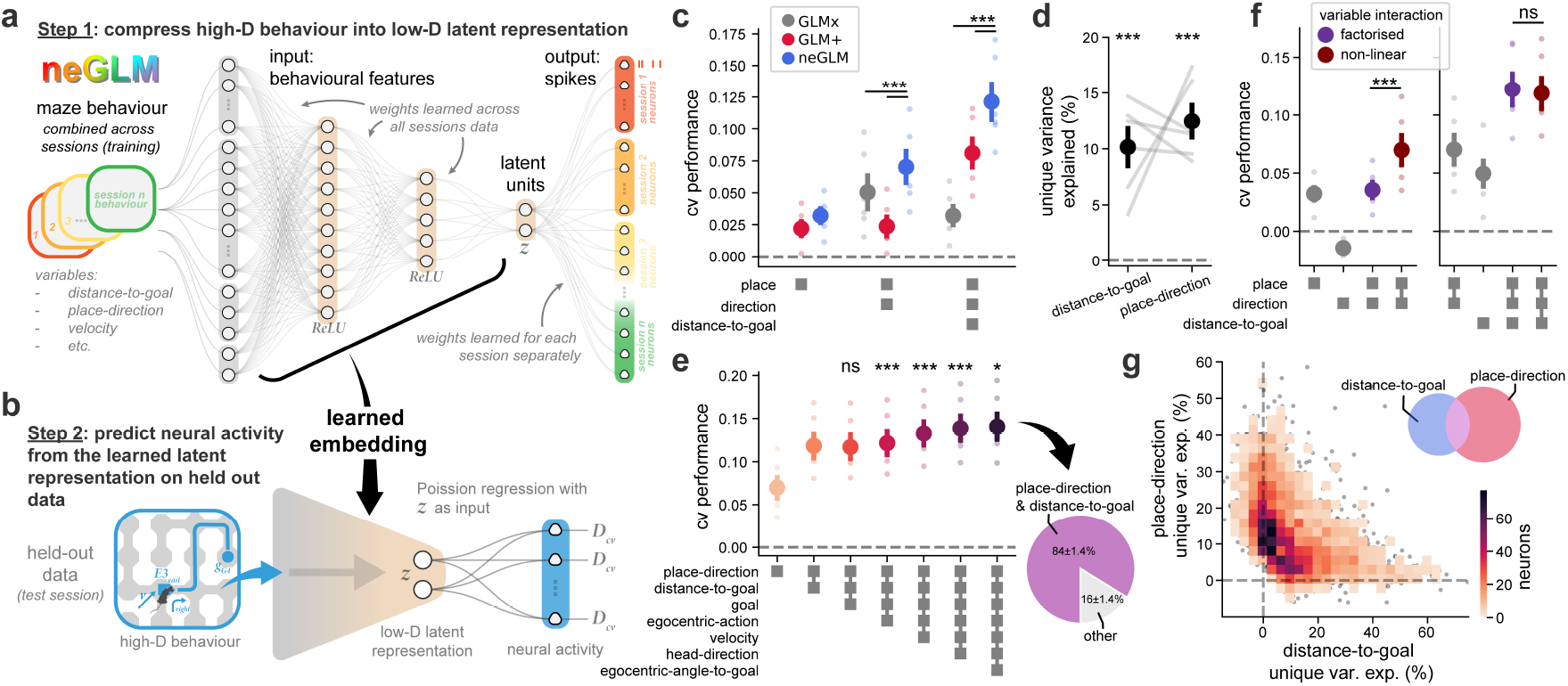
Factorised representations of place-direction and distance to goal in mFC. **(a-b)** Neural Embedding GLM (neGLM): model architecture with schematic of training and test procedures. **(c)** neGLM performance (blue) compared to baseline GLMs (GLM+ (red): Poisson GLM w/ separate one-hot encoded variables, GLM× (grey): Poisson GLM w/ single one-hot over the full input feature product space) under increasing numbers of input variables (place, direction, distance-to-goal). **(d)** Both place-direction and distance-to-goal explain unique variance in mFC. Unique variance explained determined by comparing performance of full (place-direction & distance-to-goal) and reduced (place-direction *or* distance-to-goal) neGLMs. Plotted: average unique variance explained across all cells. **(e)** Place-direction and distance-to-goal explain the majority of variance in mFC. Plotted: performance of neGLM models with increasingly large numbers of input features. Small amounts of additional variance were explained by: egocentric actions, movement velocity, head-direction, and egocentric angle to goal. Right: pie chart illustrating fraction of total explainable variance captured by place-direction & distance-to-goal. **(f)** Comparing neGLMs with different embedding architectures (see Figure S9a) can arbitrate between non-linear (maroon) and factorised (purple) interactions between represented variables. These comparisons reveal that place and direction interact non-linearly (left: performance of non-linear neGLM outperforms factorised neGLM) and that place-direction and distance-to-goal make factorised interactions (right: performance between non-linear and factorised models was not different across subjects). Bottom: Input variable interaction key, squares represent which features are present in the neGLM model, lines connecting squares indicate which variables were embedded in the same partition. **(g)** Place-direction and distance-to-goal were represented by largely distinct neural populations. Plotted: unique variance explained by place-direction and distance to goal across cells as a 2D histogram. Inset: Venn diagram of the significant representational overlaps across cells. All panels: error bars represent SEM across subjects. Grey lines and coloured dots represent individual subject values. Paired, or one-sample, *t*-tests *>* 0: *p<.05, **p<.01, ***p<.001.

We validated our approach by comparing neGLM performance to baseline GLMs with increasing numbers of input variables (place, direction, and distance-to-goal: Figure 6c). The baseline GLMs used Poisson regression to predict spikes directly from the input features, where inputs were represented as either separate one-hot vectors for different features (GLM+, Figure 6c), or as a single one-hot vector over their full product space (GLM×, Figure 6c). These two baselines represent different trade-offs between complexity and expressivity. GLM+ reduces the input dimensionality by ignoring interactions between variables, while GLM× allows for arbitrary non-linear interactions at the cost of a combinatorial explosion in features. The neGLM model performed significantly better than both baselines (paired *t*-tests, BH-corrected: *T* (5) *>* 8.9, *p <* .001), where the performance increase was largest when the input feature space was large.

Putting neGLM to work, we compared models with both place-direction and distance-to-goal inputs against models lacking one or the other, finding that both explained unique variance (Figure 6d; *t*-tests *>* 0: place-direction: *T* (5) = 4.85, *p* = .002, distance-to-goal: *T* (5) = 6.62, *p <* .001). When running neGLM models with increasing numbers of behavioural input features (added in order, Figure 6e), place-direction and distance-to-goal captured the majority of explainable variance across subjects (84±1.4%). Egocentric actions, velocity, head-direction, and egocentric-angle-to-goal each explained a small amount of additional variance (Figure 6e; paired *t*-tests rel. to next largest model, BH-corrected: +goal: *T* (5) =− 1.9, *p* = .94, +egocentric-action: *T* (5) = 7.4, *p <* .001, +velocity: *T* (5) = 15.5, *p <* .001, +head-direction: *T* (5) = 7.4, *p <* .001, +egocentric-angle-to-goal: *T* (5) = 3.1, *p* = .017). Further analyses of these additional variables are shown in Figure S10 (egocentric action), Figure S11 (velocity), and Figure S12 (egocentric angle to goal).

neGLMs also enable us to characterise interactions between behavioural features. Specifically, we can compare (i) a fully flexible non-linear embedding, with (ii) a model that partitions each set of input variables into separate embeddings to enforce factorised interactions between them. When encoding just place and direction, the flexible non-linear neGLM outperformed a model where place and direction were factorised (Figure 6f –left; paired *t*-test: *T* (5) = 5.31, *p* = .003). Thus, confirming our intuition that mFC spatial tuning spanned the product space of place and direction (Figure 4). However, when encoding place, direction, and distance-to-goal, neGLMs that factorised distance-to-goal from place & direction performed as well as models that allowed non-linear interaction between all three variables (Figure 6f – right; paired *t*-tests: *T* (5) = 1.42, *p* = .215), indicating that place-direction and distance-to-goal have factorised representations in mFC.

Finally, we examined how representations of place-direction and distance-to-goal were distributed across neurons. For each cell, we computed the unique variance explained by each variable, and found that representations were strongly axis-aligned: most cells contributed unique variance to one or other variable, but not both (Figure 6g). We confirmed that the distribution of unique variance across neurons was more specialised than expected in a mixed coding regime (Figure S9b-e; permutation test: p < .001). Together, these results demonstrate that place-direction and distance-to-goal are represented separately in mFC. Next we explore how flexible interactions between these factorised codes could implement decision-related computations.

### Offset theta-modulated codes for place-direction and distance-to-goal

We have shown that mFC activity contains a putative evaluation signal — the distance-to-goal — and a structured place-direction representation of animals’ current behaviour. Finally, we explore how these representations could interact within LFP theta-cycles to compute goal-directed decisions.

mFC and the hippocampal formation are functionally coupled via theta oscillations [44–47]. Hippocampal place cells and entorhinal grid cells encode animals’ position as they navigate [33, 48]. Within individual theta cycles, these representations sweep ahead of the animal, alternately sampling possible futures at decision-points [18, 49, 50, 17]. We hypothesised that the mFC distance code might *evaluate* these possible futures, to *update* a policy representation over successive theta cycles.

We observed a clear theta peak in the LFP spectrum during navigation (Figure 7a-b, Figure 3g) and elevated theta power relative to other trial phases (Figure 3h, Figure S13a). Theta power ramped as animals approached decision points, and was significantly elevated before decisions where subjects chose optimal actions over suboptimal habitual alternatives (Figure 7c; paired *t*-tests, BH-corrected (0.1-0.3 s pre decision; *T* (5) = 2.80, *p* = .039). This effect persisted after controlling for speed and distance-to-goal (Figure 7d, Figure S13b; *t*-test *>* 0, habit vs. structure regressor: *T* (5) = 2.91, *p* = .016). Notably, mFC silencing most impaired performance on precisely these decisions (Figure 2e), linking theta-modulated mFC dynamics to mFC-critical decisions that require knowledge of maze structure.

**Figure 7:**
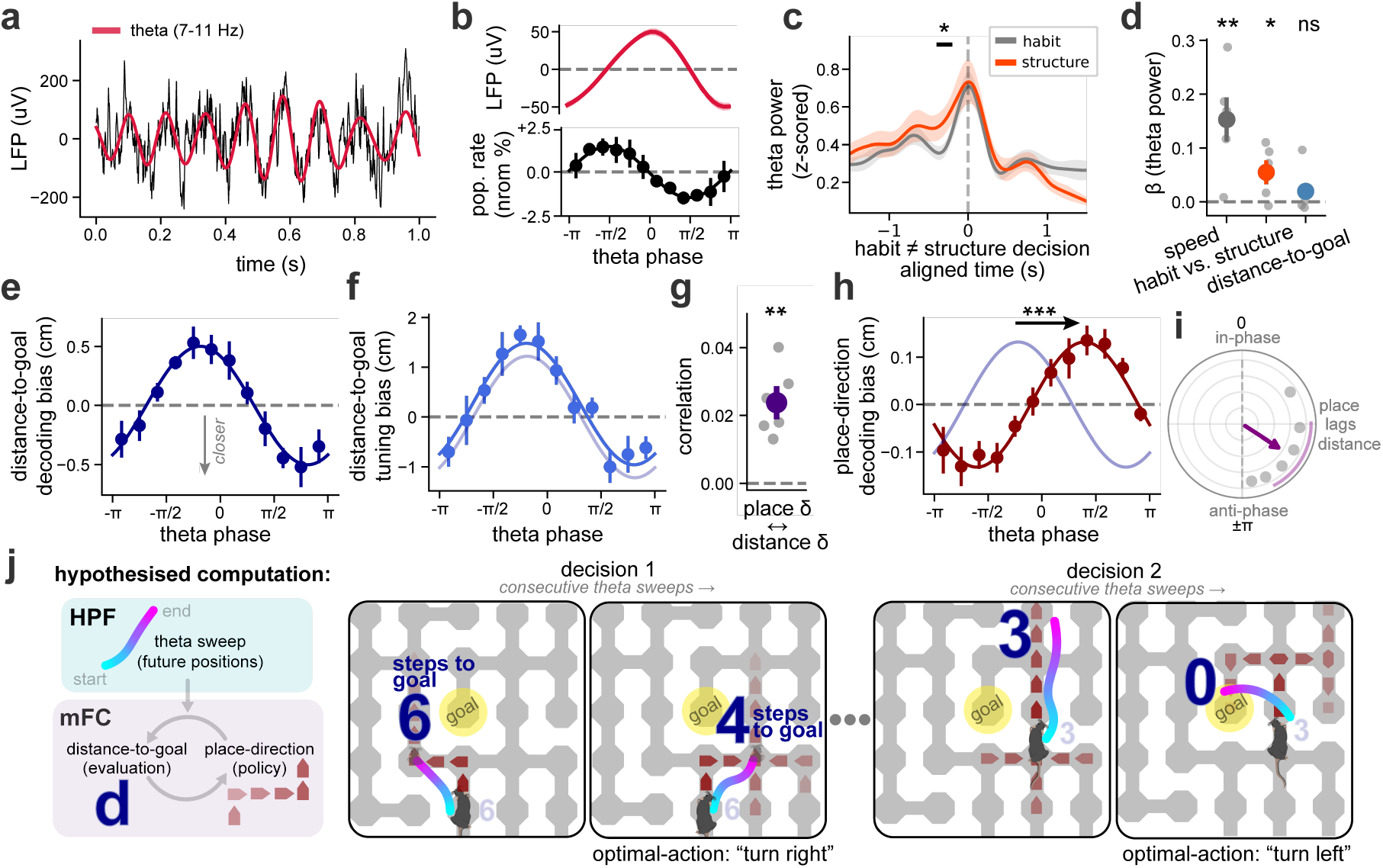
Theta modulated representations in mFC. **(a)** Example local-field potential (LFP) signal with band-passed theta component. **(b)** Average LFP signal (top) and population activity (bottom) aligned to LFP theta-phase. Bottom: average modulation profile across subjects (black markers) fit with a sinusoid (full line). **(c)** Average theta-band power, aligned to decisions where structure-based and habitual strategies prescribed different actions. Theta ramped more strongly before structure-based (optimal; orange) than habitual (grey) choices. Statistical test: paired *t*-test per time-point (BH-corrected) **(d)** Choosing the structure-based action predicted higher theta power independently of running speed and distance-to-goal, shown as per-subject regression coefficients (*β*) for theta power in the pre-decision window identified in c. **(e)** The distance-to-goal representation processed from further to closer to goal within a single theta cycle. Plotted: Average distance-to-goal decoding bias as a function of theta phase. Negative values represent decoded distances that were closer to the goal than average. Formatted as in panel c. **(f)** Confirmation of theta-modulation in the distance code. Plotted: the necessary shift (in distance-to-goal) required to align distance-to-goal tuning curves derived from spikes at different theta phase to a reference average. Formatted as in panel c, with reference fitted sinusoid from e in faint navy. **(g)** Distance-to-goal and place-direction representations fluctuate together on a behavioural timescale. Plotted: correlation of distance-bias and position-on-trajectory bias derived from independent decoders. Statistical test: one-sample *t*-test *>* 0 across subjects. **(h)** The place-direction representation was theta modulated and systematically offset from the distance-to-goal representation. Plotted: average bias of a place-on-trajectory decoder as a function of theta-phase. Formatted as in panel f. Statistical test: subject-level bootstrap of phase offset difference. **(i)** Phase offset between place-direction and distance-to-goal theta modulation profiles. Plotted: purple arrow represents the phase-offset circular mean across-subject; shaded arc represents bootstrap 95% CI on the mean. **(j)** Hypothesised computation: hippocampal theta sweeps sample possible future locations at decision points (cyan-to-pink lines), mFC reads out their distance-to-goal, and the evaluated samples update a policy represented by the place-direction code. All panels: error bars and shaded errors represent SEM across subjects. Individual subject values plotted as faint grey lines or dots. Markers for statistical tests: *p<.05, **p<.01, ***p<.001

We reasoned that if the mFC distance code evaluates possible futures sampled by hippocampal theta sweeps, the distance-to-goal signal should be modulated by theta phase. To test this, we split mFC activity into twelve theta phase bins, trained a decoder to predict distance-to-goal from the average neural activity across phase bins, then tested the decoder on spikes from each theta phase separately (Figure S13d,f). This revealed a systematic bias in distance-to-goal representations across theta phases: mFC represented shorter distances at the trough of theta than at its peak (Hotelling’s *T* ^2^ test: *F* (2, 9) = 7.35, *p* = .013; Figure 7e). A separate analysis, testing whether distance tuning curves were shifted for spikes occurring at different theta phases, recovered the same effect (Hotelling’s *T* ^2^ test: *F* (2, 9) = 15.46, *p* = .001; Figure 7f, Figure S13g). Notably, this modulation profile was distinct from the average theta modulation of population firing rates (Figure 7b - bottom, Figure S13c).

If the place-direction code is part of the same theta-paced computation, both codes should be coupled dynamically. To test for behavioural-timescale coupling, we trained independent decoders for place-direction and distance-to-goal on disjoint neural populations — cells with unique variance explained by only one of the two variables. For each decoder, we computed a ‘directional bias’ per 200-ms bin: the distance-weighted sum of posterior probabilities at previous, current and next locations/distances on a linearised trajectory, indicating whether represented locations/distances were ahead of or behind the animal. The two biases were significantly correlated (Figure 7g; mean Pearson *r* = .024± .004 across subjects, *t*-test *>* 0, *T* (5) = 5.8, *p* = .001), indicating that place-direction and distance-to-goal codes fluctuated together on a behavioural timescale.

To examine theta-timescale coupling we analysed how the place-direction code varied within theta cycles. We did not record sufficient neurons and behaviour to reliably decode trajectories within individual theta sweeps; instead, we applied the same logistic regression decoder to predict subject’s position on trajectory from theta-stratified mFC activity, and computed the average directional bias at each theta phase. This revealed significant theta-modulation of the place-direction representation (Hotelling’s *T* ^2^ test: *F* (2, 9) = 10.57, *p* = .004, Figure 7h). Strikingly, the theta modulation of the place-direction code was delayed by 1/3 of a cycle (∼40 ms) from that of the distance-to-goal code (Figure 7i; 2.214 rad.; 95% *CI* [1.53, 2.67], *p <* .001; Rayleigh test: *Z*(5) = 3.49, *p* = .016).

Thus, the place-direction and distance-to-goal codes are coupled — both on a behavioural timescale and within individual theta cycles, where the distance code sweeps from far to near the goal, and the place-direction code follows. These dynamics suggest a mechanism for goal-directed decisions: hippocampal theta sweeps sample possible futures; mFC evaluates each by its distance-to-goal; and the resulting evaluation pushes the place-direction policy towards optimal actions —– those that decrease distance-to-goal (Figure 7j).

## Discussion

Our data causally implicate mFC in flexible goal-directed navigation (Figure 2), and provide a detailed characterisation of the underlying neural tuning and dynamics. Neural activity was dominated by two factorised representations: an efficient embedding of behavioural trajectories (Figure 4), and a population code for shortest-path distance-to-goal (Figure 5). Both were theta-modulated and offset within the cycle, suggesting a computation in which mFC evaluates possible futures sampled by the hippocampus to update a structured policy representation (Figure 7).

Our optogenetic results establish that mFC is required for flexible navigation in complex environments, even when goals are fully observable. This contrasts with cued water-maze navigation, where mFC lesions leave performance largely intact [51, 27]. The deficit was present during expert behaviour, in contrast to spatial alternation, reversal, and outcome devaluation tasks, where mFC-dependence is confined to learning [30, 28, 27, 29, 8]. Together, these results suggest that mFC is critical not only during learning, but also when knowledge of environment structure must be used to flexibly select goal-directed actions. mFC inhibition caused subjects to default to habitual choices and fail to course-correct after errors — reminiscent of patients with prefrontal damage, who similarly show perseveration and impaired flexibility in novel situations^1^ [1–3, 52, 4, 53].

A subpopulation of mFC neurons were tuned to conjunctions of maze location and movement direction, with low-dimensional tuning that reflected both the structure of the environment and the statistics of behaviour. This code may represent the subject’s behavioural policy, a model of the environment’s structure, or both. Consistent with the former role, place-direction tuning formed an efficient basis set for representing navigational trajectories (Figure 4m), and predicted future choices (Figure 4n). These findings extend efficient-coding principles from sensory cortex [35, 54–56] to representations of behaviour, echoing proposals that the brain compresses behavioural policies [57]. The structure of place-direction tuning also suggests a hierarchical organisation of behaviour [58], with many tuning patterns interpretable as abstract or temporally extended actions (e.g., corridor, subgoal, exit dead-end). Efficient coding and hierarchical organisation may reflect a common process, as compression of behavioural sequences is a possible mechanism for discovering useful macro-actions [59, 60].

A second subpopulation of mFC neurons formed a population code for shortest-path distance to goal, with different neurons preferring different distances. Distance signals have been reported in single neurons across the brain [61–63] and in human fMRI [64, 65] but not, to our knowledge, as a population code tiling shortest-path distance. Our finding also differs from “progress-to-goal” coding in frontal cortex during navigation [66, 38, 67] and timing tasks [68], perhaps because our mice cannot predict paths from one trial to the next. In our task, shortest-path distance-to-goal corresponds to goal-conditioned state value, connecting our findings to value coding in frontal cortex [69, 70] and to evidence that internal value signals control behaviour [71–73].

mFC and the hippocampal formation are functionally coupled through theta oscillations [74–76], with coupling modulated by task demands [77] and causally important for navigation and value-learning behaviours [72, 78]. Co-recordings during navigation have revealed structured coordination, with hippocampal spatial sweeps aligning to mFC representations of reward expectancy [44] or upcoming choices and paths [46]. However, linking these observations to a concrete decision mechanism has been hampered by the representational ambiguity of mFC in simple tasks.

Our finding of co-theta-modulated but temporally offset codes for distance-to-goal and place-direction suggest a possible mechanism, characterised by a *sample—evaluate—update* loop operating at theta frequency: hippocampus samples possible future locations, these are evaluated by the distance-to-goal code, with evaluated samples used to update a policy represented by the place-direction code. This interpretation aligns with recent proposals that sequential activity in the hippocampus updates policy representations in prefrontal cortex [79, 80]. Further, recent observations that hippocampal theta sweeps are themselves goal- and attention-directed [81, 49] raise the possibility that updated mFC policies bias subsequent hippocampal sampling towards more promising futures — yielding an iterative policy refinement process [79].

This account remains provisional in two respects. First, we recover only phase-averaged modulation profiles for the two codes: their significant offset shows that the two sweeps occupy distinct phases of the theta cycle, but averaging across cycles cannot establish which code drives the other from one cycle to the next. Second, the proposed algorithm pre-supposes a distance-to-goal computation, yet how this quantity is derived remains unclear. While information about the subject’s current position likely reaches mFC from the hippocampal formation [14– 16], information about the current goal could come from another frontal region [38], the hippocampal input to mFC [82], or somewhere else entirely. Resolving these ambiguities will require dense, simultaneous recordings across the hippocampal formation and frontal cortex, together with closed-loop perturbations that probe the causal flow of information across the two regions.

## Author contributions

CRediT [83] author contribution statement. **PTD**: conceptualisation, software, methodology, validation, formal analysis, investigation, data curation, writing (original draft & revisions), visualisation. **KTJ**: software, formal analysis, writing (review & editing), supervision. **YC**: investigation (animal experiments). **CDGB**: software, methodology. **CQ**: conceptualisation. **BSG**: conceptualisation, methodology. **JLE**: investigation (animal experiments), **RJC**: investigation (histology), **MEW**: conceptualisation, supervision, funding acquisition. **TEJB**: conceptualisation, formal analysis, resources, writing (review & editing), supervision, project administration, funding acquisition. **TEA**: conceptualisation, methodology, software, formal analysis, investigation, resources, writing (review & editing), supervision, project administration, funding acquisition.

## Data & code availability

Datasets (ephys & opto experiments) have been deposited on Zenodo. Each is accompanied by code repositories for regenerating analyses in this manuscript (analysis code) and quick-start tutorials for setting up new analyses projects (tutorial code). Please note data, code and associated documentation are subject to change before final publication of the manuscript.

- Ephys dataset:
  – data
  – analysis code
  – tutorial code
- Opto dataset:
  – data
  – analysis code
  – tutorial code
- Maze Design

## Acknowledgments

We are grateful to Laurence Hunt, Sebastijan Veselic & Neil Burgess for their feedback on the manuscript. This work was supported by the Oxford Clarendon Fund (PTD); a Wellcome Trust Career Development Award (225926/Z/22/Z; TA); a Wellcome Principal Research Fellowship (219525/Z/19/Z; TEJB, KTJ); the Gatsby Initiative for Brain Development and Psychiatry (GAT3955; TEJB); the Jean Francois and Marie-Laure de Clermont Tonerre Foundation (TEJB). KTJ & TEJB were also supported by the Sainsbury Wellcome Centre’s core provided by Wellcome (219627/Z/19/Z) and the Gatsby Charitable Foundation (GAT3755). We are also grateful to the Sainsbury Wellcome Center IT and histology teams for their ongoing support.

## Methods

### Behavioural apparatus & task

Experiments were performed on a custom-built maze apparatus [25] controlled with pyControl [84] (Figure S1a). Pretraining was performed on a 3-by-3 maze and the main experiment and recordings were performed on a 7-by-7 maze. Each maze consisted of towers organised in a grid, each with a stimulus LED, reward port with infrared beam and solenoid valve for dispensing liquid rewards, and speaker, interconnected by removable walkways to create complex maze layouts. On each trial, a randomly chosen goal (tower) was cued by illuminating its LED and playing a brief sound (white noise, 0.5 s). Subjects then navigated to the cued goal and received a water reward, followed by a 4-8 s inter-trial interval (ITI) (Figure 1a). Maze design specifications can be found here.

### Maze design

Optimised mazes were generated using a custom designed genetic algorithm developed in Python with NetworkX [85] (Figure S1c). First, 100 random mazes were generated that satisfied the following criteria: mazes defined a connected graph (no disconnected nodes), the total number of edges was less than 46, that the total number of outer edges (on outer perimeter) was less than 19, that all open connected components (Figure S1c - bottom right) were less than size 3 and that all closed connected components (Figure S1c) were less than size 8. The initialised maze with the highest maze fitness was kept for further optimisation, where maze fitness was defined as:

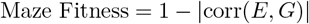

Where, *E* and *G* are the sets of all Euclidean and shortest-path (geodesic) distances between all pairs of towers on the maze, respectively. Next, all one-step variants of this top-ranked maze were generated (i.e. all permutations of the maze with one edge added or removed - Figure S1c). The resultant mutant with the highest maze fitness, that also satisfied the above criteria, was retained. Subsequently all one-step variants of this maze were generated, and this process was repeated until the maze fitness values converged or some maximum number of mutation cycles was reached, yielding an “optimised” maze.

### Goal selection

For the maze optogenetic experiment all 49 maze towers were used as goal locations (for both Mazes 1 & 2). For the maze electrophysiology experiment, 24 goal locations were sub-selected, and on some later sessions (after learning) further 12 goal subsets were used to increase the number of trials per-goal to aid goal decoding analyses. Goal subsets were selected with another custom genetic algorithm. First, 24 towers were randomly selected as goal nodes on an optimised maze. Next, the “goal fitness” was calculated for each goal node and non-goal node on the maze, where goal fitness was defined as:

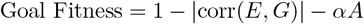

Where, *A* was the number of goal locations immediately adjacent to the selected node, weighted by some factor *α* (0.15). The adjacency penalty was introduced to avoid clustering of goal locations and encourage exploration of the entire maze. Next, the goal location with the lowest goal fitness was replaced with the non-goal location with the highest goal fitness. These steps were repeated until the average goal fitness across the goal set converged, or some maximum number of permutations was reached, yielding an “optimised” goal-set on a given maze. Goal locations were optimised as above for Maze 1 and then held constant for Maze 2 and the Rooms Maze. Further, some sessions with 12-goals (subset 1 & subset 2) were used on late sessions, these were chosen by hand (Figure S1b).

### Subjects

A total of 26 male and female C57BL/6J mice (6 weeks old at start of experiments) were used. Animals were group-housed before surgery, on a 12 h light/dark cycle. Animals that were implanted with silicon probes were individually housed after surgery. For the maze optogenetic experiment, animals (M:8, F:8) first underwent viral injections and optic fibre cannulae implantation before pretraining and maze experiments. One subject had post-surgical complications and did not participate in the experiment. For the stGtACR2 electrophysiology experiments, animals (M:2) first underwent surgery and then recordings 1-2 times a week for 6 weeks. For the maze electrophysiology experiment, animals (M:8) were first pretrained then the best performing six subjects were implanted with silicon probes and performed maze experiments. Sample sizes were not predetermined statistically but were comparable to previous studies [86, 38]. All procedures complied with Oxford University guidelines and were conducted under UK Home Office Project Licence PP6834709.

### Maze behavioural training

Subjects were placed on water restriction 24 hours before starting behavioural training and were trained for no more than 14 consecutive days without a day off, where they received 1 hour ad libitum water access in their home cages. On training days, mice typically received all their water from the task but were given additional water if required to maintain their body weight above 80% of their pre-restriction baseline weight.

Before exposure to the large 7-by-7 mazes, subjects underwent a standardised pre-training protocol on smaller 3-by-3 small mazes (Figure S1d). Subjects were first exposed to the maze apparatus together in cage groups for 15 minutes. Next, subjects underwent magazine training to discover the reward ports on each tower. Here all towers were cued with the stimulus LED until subjects poked into a given reward port, water rewards were delivered and the light cue was extinguished. The session ended after all rewards were collected. Next subjects were exposed to the full task on a series of structured mazes over multiple sessions. Reward sizes were initially very large and ITI periods very long to make cue and reward events very salient. With training reward-sizes reduced and ITI periods were shortened, increasing the number of trials completed in a single session to ∼100 (Figure S1e) over one week. Further details on the pretraining procedure can be found here.

For maze optogenetic experiments, the cohort was split into two groups with mice experiencing either Maze 1 then Maze 2 or Maze 2 then Maze 1 (randomised and counterbalanced across opto and control conditions). Mice learned the 1^*st*^ Maze for 8 days before optogenetic manipulations over expert behaviour (13 days), mice then learned the 2^*nd*^ Maze for 5 days before 12 further days with optogenetic manipulations. For maze electrophysiology experiments subjects were exposed to Maze 1 for 13 days, then Maze 2 and the Rooms Maze for 11 days.

### Surgery

Subjects were taken off water restriction 72 hours prior to surgery. During surgery subjects were anaesthetised with inhaled isoflurane (3% induction, 0.5-1% maintenance). They were then treated with buprenorphine (0.1 mg/kg, sc), meloxicam (5 mg/kg, sc), bupivacaine (8 mg/kg, sc) and a sodium chloride (0.9% w/v) and glucose (5% w/v) solution (40 *µ*L/g, sc), before being placed in a stereotactic frame (Kopf Instruments, USA) under an OPMI Lumera S7 surgical microscope (Carl Zeiss AG, DE). Mice were maintained at 37 ^◦^C using a rectal probe and heating blanket (Harvard Apparatus). Mice were allowed to recover from surgery for a minimum of 7 days, being closely monitored and given additional doses of meloxicam (5 mg/kg, p.o; formulated as jelly) each day over the first 3 days post-surgery.

For maze optogenetic experiments two 0.6 × 0.6 mm craniotomies were made over mFC, then 600 nL of AAV1-CKIIa-stGtACR2-FusionRed (diluted to 1 × 10^12^ vg/mL, addgene #105669, opto group) or AAV1-CamKIIa-mCherry (diluted 1 × 10^12^ vg/mL, addgene #114469, control group) was injected into each hemisphere targeting the prelimbic subregion of mFC (AP: 2.0 mm, ML: -0.4 mm, DV: -1.2 mm from surface). Next, two 200-*µ*m-diameter ceramic optical fibres were implanted chronically targeting the injection sites at a 10^◦^ angle.

For maze electrophysiology experiments, a ground screw was implanted over posterior lateral cortex, and a 1 × 1.5 mm craniotomy was made over mFC. Next, a preassembled implant was lowered into position with the silicon probe tip targeting the prelimbic region of the left hemisphere (AP: 2.0 mm, ML: -0.4 mm, DV: -1.0 mm from surface). The implant consisted of a 6-shank Cambridge Neurotech F-series silicon probe (Cambridge Neurotech, UK) mounted to a Nanodrive (Ronal Tool Inc, York, PA, USA) and connected to a Mini-amp-64 amplifier (Cambridge Neurotech), all housed within a custom 3D printed plastic enclosure attached to a 3D printed steel base-plate (Figure S4a). The craniotomy was sealed with Dura-gelTM (Cambridge Neurotech, UK) and the base-plate of the implant was securely attached to the subjects’ skull with self-adhesive resin cement (RelyXTM Unicem2 automix; 3M, USA).

For stGtACR2 electrophysiology experiments, 500 nL of AAV1-CKIIa-stGtACR2-FusionRed was injected into the left mFC (as above) then a pre-assembled implant containing a 200-*µ*m-diameter optic-fibre and Neuropixel 1.0 probe (Imec) were implanted chronically at the same site (fibre tip targeted to AP: 2.0 mm, ML: -0.4 mm, DV: -1.0 mm from surface). Ground screws implanted, craniotomies sealed, and cemented as above.

### Histology

At the end of each experiment, silicon probes were recovered under terminal pentobarbital anaesthesia (200 mg/kg, i.p.). Once anaesthetised, mice were secured in a stereotactic frame and the implant body was detached from the base-plate. Whole brains were then harvested from each animal and fixed in paraformaldehyde for histological preparation. For electrophysiology experiments brains were sliced at 50 *µ*m, slide mounted and Nissl stained before automated bright field imaging (Zeiss Axioscan-Slidescanner). For optogenetic experiments fixed whole brains were imaged with serial 2p tomography as described previously [87].

### Optogenetic silencing

Once mice had learned a given maze structure, on a random 25% of navigation trials mFC was inhibited bilaterally (light-ON trial) by delivering blue light via a 427 nm LED (Doric) slip-ring bearing mounted to a custom 200-*µ*m 0.66-NA optic fibre patch cords (Doric) tethered to the animal. During light-ON trials light was delivered as a square-wave pulse at ∼ 1.0 mW optical power at the fibre tip, 1-3 s before the goal was cued up until subjects reach the goal and received a reward, or a max of 30 s. This manipulation was made over 12-13 days of stable behaviour on each maze. It took control subjects ∼5 days to adjust navigating with the light during light-ON trials (Figure S3c), thus, for all summary analyses trials from these first 5 days on the opto protocol were excluded (from both groups).

### stGtACR2 validation

To validate our optogenetic approach we implanted two mice with Neuropixel 1.0 probes and optic fibres, and expressed the stGtACR2 construct in mFC. Electrophysiology data was collected via PXIe 1000 acquisition module (Imec) mounted on a National Instruments (NI) PXIe-1090 chassis using OpenEphys software, with blue light (427 nm, 0.5 mW or 5 mW) delivered through a tethered optic fibre in the animal’s homecage. Recording sessions consisted of continuous ephys recordings with randomly spaced light pulses (1, 10 or 30 s) over ∼ 40 min sessions. From these recordings we quantified the efficacy of neuronal inhibition over a 6 week period as well as minor off-target effects (Figure S2e-j).

### Locomotor controls for optogenetic manipulations

To quantify the baseline effects of mFC inhibition on spontaneous locomotion, subjects from the maze optogenetic experiment were also exposed to a 1 m × 1 m open-field arena for 30 min after some maze behaviour sessions. The open-field was positioned inside the maze arena (dark) where mice foraged for randomly scattered chocolate sprinkles (DeRuijter), while blue light (427 nm, 1.0 mW) was delivered to mFC via a tethered optic fibre for 10 s every 30-90 s with concurrent video recordings. From these recordings we quantified locomotion statistics aligned to light-ON and light-OFF events (Figure S2k-m).

### Chronic electrophysiology

For maze electrophysiology experiments using Cambridge Neurotech probes, neural activity was recorded at 30 Hz with a 64-channel Miniamp-64 amplifier board (hardware bandpass filtering between 1.1 and 7603.8 Hz; Cambridge Neurotech) connected to an OpenEphys acquisition board via an ultra-thin flexible serial peripheral interface cable (Intan technologies, USA) and a custom-built commutator. The commutator functioned to prevent tangling of the flex-cable during freely moving behaviour. Electrophysiological and behavioural data were synchronised by sending sync pulses from the pyControl system to the OpenEphys acquisition board.

Probes were advanced ventrally in 150 *µ*m increments throughout the experiment to a final depth of -1.6 mm from the cortical surface. Specifically, probes were advanced 3 days prior to the main experiment (experimental day -3) and 4 days before subjects finished/changed maze configurations (experimental days 8, 20 and 32). This was to increase the number of neurons sampled throughout the experiment and to buffer against potential reductions in neuronal activity over the recording period, while ensuring that the same groups of neurons could be compared both within and across mazes.

### Electrophysiology preprocessing

Raw electrophysiological data was preprocessed and spike-sorted according to IBL recommendations [88], in a custom pipeline built with SpikeInterface [89]. First raw data was preprocessed by automatically removing dead/bad channels and applying common average referencing per shank to the signal on the remaining channels. Preprocessed data was then spike-sorted with Kilosort 4 [90] where *Th*_*universal*_ and *Th*_*learned*_ parameters were set to 11 and 10, respectively. We then derived a variety of quality metrics for each putative cluster from the spike-sorting output (ISI violations ratio, amplitude cutoff, presence ratio, median amplitude, SD ratio and average firing rate; see SpikeInterface documentation for further details). “Single units” were defined as spike-sorted clusters with: ISI violations ratio < 0.1, amplitude cutoff < 0.1, average firing rate > 0.1 Hz, presence ratio > 0.95, median amplitude > 30 *µ*V, SD ratio > 3. “Multi-unit activity” was defined as spike-sorted clusters that were not single units and had: ISI violations ratio < 0.6, median amplitude > 20 *µ*V, presence ratio > 0.9.

### Video preprocessing

Video recordings of maze and open-field behaviour were captured at 60 Hz with 1280×1024 resolution using a Chameleon3 camera (CM3-U3-13Y3M; Teledyne FLIR, USA) with wide-field lens at 2.8 mm focal length (Computar 2.8-12 mm 1:1.3, IR 1/3”). Markerless pose estimation was performed using DeepLabCut (maze electrophysiology experiments) [91] or SLEAP (maze optogenetics experiments) [92]. DeepLabCut/SLEAP outputs were further processed to remove tracking errors, and map from video pixels to physical coordinates, with custom Python code.

### Histological preprocessing

For maze optogenetic experiments, brains (imaged with serial 2p tomography) were automatically registered to the Allen CCFv3 atlas (10 *µ*m) [93] using brainreg [94]. The entry points and tips of subject’s optic fibres were manually annotated in atlas space using the BrainGlobe Napari viewer [94]. The total volume, and corresponding anatomical regions, of mFC inhibition in each subject were estimated as the union of two cones extending fibre tips (length: 500 *µ*m, half-angle: 35^◦^) and thresholded viral expression in atlas space, using custom Python code, visualised with brainreg [94] (Figure S2b-d).

For maze electrophysiology experiments, probe tracts (from single slice histology) were first manually identified and ported into HERBS [95], slices were then individually registered to Allen CCFv3 (25 *µ*m) atlas. Between 2 and 6 shanks could be identified in each subject. We then found the plane of best fit in CCFv3 space for each subject’s reconstructed tracts and determined consensus probe positions by evenly spacing six shanks across this plane (inferring the position of any missing shanks). Final probe depths could not easily be determined from histology and were instead taken as initial depth of the probe noted during implantation, plus any additional distance from advancing the microdrive. Contact positions and anatomical regions were determined relative to this probe depth. Further, spike-sorted clusters were assigned to the anatomical region of the contact with the largest template waveform.

### Maze navigation metrics

“Excess steps” were calculated per trial as the difference between the length (in tower to tower steps) of the subject’s real trajectory and the optimal shortest-path length from the start location at cue to the goal location. “Backtracks” refer to any time a subject returned to a tower just visited. “Navigational errors” refer to any time subjects took an action that increased shortest-path distance-to-goal after it was previously decreasing. “Repeat errors” refer to any “navigational errors” that occurred at the same location on the same trial more than once.

Linear mixed-effects models with subject included as a random intercept were fitted using maximum likelihood to test the effects of day on maze and the maze order × day on maze interaction on excess steps and trials completed per session across optogenetic and electrophysiology experiments. A mixed-design ANOVA with light-ON trial as a within-subject factor and group (opto or control) as a between-subject factor was conducted to test for a significant group × light-ON interaction on excess steps and other performance metrics in the optogenetic experiment.

### Mixture-of-strategies modelling

At every navigational decision (i.e. each tower during the navigation period), we assigned each option a value given by a weighted linear combination of four components, each with its own weight:

- a vector-navigation component (*Q*_*v*_, weight *w*_*v*_), the cosine of the angle to the goal, favouring options that move towards it;
- a structure-based component (*Q*_*s*_, weight *w*_*s*_), set to 1 for the option(s) reducing the shortest-path distance to the goal and 0 otherwise;
- a habit component (*Q*_*h*_, weight *w*_*h*_), the probability of an action given the subject’s current and previous position, independent of the current goal (see above); and
- a backtracking penalty (*Q*_*p*_, weight *w*_*p*_), set to −1 for options returning the subject to the just-visited location.

These values then determined choice probabilities via a softmax function:

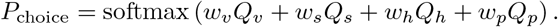

In both experiments we fit the component weights by maximum likelihood, pooling choices at the level appropriate to each analysis. For the electrophysiology experiment, we fit weights separately for each subject – pooling choices across that subject’s late sessions (the last 7 days on each maze) – and, separately, for each maze-day, pooling across all subjects. For the optogenetic experiment, we fit weights separately for each subject and light condition (light-ON vs light-OFF). To test whether a strategy contributed to subjects’ navigational decisions, we performed a one-sample *t*-test against zero (*>* 0) on its weights across subjects. To compare weights across conditions and trial types (light-ON/light-OFF) in the optogenetic experiment, we entered them into a mixed ANOVA (as described above).

### Navigation strategies comparisons

To quantify the relative contribution of each strategy to behaviour, we measured the rate of structure-based (i.e. optimal) choices at decision points over late sessions on Maze 1 and Maze 2, where strategies were most decorrelated (over the last 7 days of behaviour on each maze). We restricted analyses to decisions where the alternative strategies prescribed suboptimal actions – that is, where the option(s) preferred by the vector strategy, the habit strategy, or both did not coincide with the optimal structure-based option. Within each strategy conflict scenario we computed the average probability each subject took the optimal structure-based action. This was compared to the chance level expected from a uniform-random choice among the available options at each tower (per subject), with structure-based choice probabilities tested against chance using paired *t*-tests across subjects.

In the optogenetic experiment we applied the same analysis, but additionally resolved each strategy comparison by experimental group (opto vs. control) and by inhibition (light-ON vs. light-OFF trials). Within each conflict scenario, the effect of mFC inhibition on the probability of optimal choice was assessed with a 2-way mixed ANOVA (group × inhibition, as above). To ask whether this effect varied with the prevailing strategy conflict, we then ran a 3-way mixed ANOVA across scenarios (group × inhibition × scenario) to test whether the inhibition-induced change in optimal choice differed between opto and control animals across strategy agreement and conflict.

### Trial-& event-aligned neural activity

We aligned activity in two ways, both yielding firing rates evaluated every 40 ms with a Gaussian kernel. For *trial alignment*, we time-warped the intervals between trial events (cue, reward, end of reward consumption, and end of the inter-trial interval) to the median of those intervals across all late sessions, leaving pre-cue activity unwarped. Spike times within each interval were mapped into this common reference frame by linear interpolation between event times; because warping alters spike density, spikes were weighted by the corresponding stretch factor before rates were computed. For *event alignment*, we instead aligned spikes directly to each trial event (cue, reward, end of reward consumption).

From these rates we derived population and single-unit responses. Trial-aligned population activity was obtained by z-scoring single-unit rates across all trials in a session and averaging across the population on each trial. Single-unit responses were the trial-averaged firing rate – either across all goal locations or stratified by goal location – and were additionally z-scored across time when shown in population heatmaps.

Unit ordering in heatmaps was cross-validated: derived from data split by trial, then applied to tuning curves computed from all trials. For trial-aligned heatmaps, we clustered the first-half tuning curves with *k*-means (*k* = 6), assigned each second-half curve to its nearest centroid (Euclidean distance), and ordered units by cluster. For event-aligned heatmaps, we ordered units by the argmax of their split-half tuning curves, separately for each trial event (Figure S4e).

### LFP & CSD

Local field potential signal (LFP) across probe channels was calculated by first low-pass (450 Hz) filtering and down-sampling (30,000 Hz to 1,500 Hz) the raw mFC wideband signal. “LFP” used for further analyses was defined as the average signal across a single row of channels with good signal on either shank 3 or 4 (whichever had the most contacts passing quality control). Current source density (“CSD”) signal used for further analysis was calculated by first averaging the LFP signal across a single row of channels (with good signal) on shanks 1, 3 and 5 or 2, 4 and 6 (*C*_1_, *C*_2_ and *C*_3_) and taking the second spatial derivative of these signals:

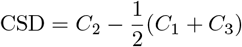

Artefacts in “LFP” and “CSD” signals were identified at times where signals deviated more than 500 *µ*V and 100 *µ*V, respectively, from the previous sample (which occurred sparingly). The signal 5 ms around these events was removed and linearly interpolated.

Using the multitaper method [96], we computed the power spectral density (PSD) of three trial phases, each trimmed by 0.5 s at both ends: navigation (cue to reward), reward consumption (reward to end of reward consumption), and the inter-trial interval (end of reward consumption to the next trial). Because these segments varied in duration, their PSDs had slightly different frequency resolutions; we therefore interpolated each onto a common, log-spaced frequency grid (1–250 Hz) and averaged across trials in late sessions for each subject. Within each frequency band of interest (theta, 7– 11 Hz; and 4–5 Hz), we compared PSD averages between trial phases using paired *t*-tests across subjects.

Time-frequency representations were computed using a continuous wavelet transform of the “LFP” or “CSD” signal. The resulting power spectrogram was then z-scored across the time dimension independently for each frequency to highlight relative changes within frequency bands across a session. These normalised spectrograms were aligned to different trial events (cue, reward, end-reward-consumption) and averaged across all trials in late sessions for all subjects, yielding the typical peri-event oscillatory power changes for each trial event.

### Allocentric goal decoding

To decode navigation goals from either trial-aligned or event-aligned neural activity, we trained either linear (multinomial logistic regression; MLR) or nonlinear classifiers (custom multi-layer perceptron classifier implemented with PyTorch) on neural firing rates calculated over 0.2 s trial-aligned windows. For session level decoding the current goal was predicted from neural activity at different time points relative to cue or reward. To combine data across multiple subjects and sessions, we constructed pseudo-population activity vectors, by combining neural activity from different navigation trials in various sessions with common alignment (e.g., timepoint) and navigational goal, yielding data matrices of pseudo-population activity by pseudotrials. Decoding analyses were performed with goal-stratified (hold one trial out per goal per session) cross-validation over a minimum of six folds, for each maze goal set separately.

To test whether allocentric goal decoding exceeded what generic spatial coding could explain, we compared two MLR decoding schemes. In the direct scheme (our baseline), goal was decoded straight from population spike counts. In the mediated scheme, we first trained an independent MLR decoder to predict a spatial variable from these spike counts – either place or place-direction – and then decoded goal from the resulting probability distribution over that variable. We compared goal-decoding accuracy across decoding schemes at time points aligned to cue, reward, or shortest-path distance to goal. For every decoder, the regularisation strength was set by an inner cross-validated search on each train–test fold.

### Distance-to-goal tuning

Distance-to-goal tuning on each navigational trial was calculated by averaging neural activity during the navigation portion of each trial, while subjects were less than 30 steps from goal (to exclude off-task activity) into equally spaced 4 cm distance bins (see below for details on distance metric definitions). Single neuron distance tuning was plotted as the mean and s.e.m. across trials, optionally normalised to the neuron’s maximum firing rate. For theta-modulation analyses (see below), tuning curves were further filtered for times when subjects were moving (speed > 0.12 m/s).

To characterise distance-tuning curves across the population we fitted each empirical tuning curve with four-parameter gamma or Gaussian fits, defined as:

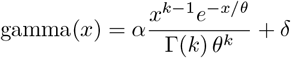

where *α, k, θ*, and *δ* are the size, shape, scale, and offset parameters, respectively, and Γ(*k*) is the gamma function.

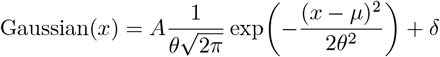

where *A, µ, θ*, and *δ* are the amplitude, mean, standard deviation, and offset parameters, respectively.

Across subjects, distance tuning curves were best fit by the gamma distribution function and as such we used these fits for further characterisation. We derived a ‘split-half correlation’ metric for the distance tuning of each cell by calculating two distance tuning curves from random split halves of trials over 100 random splits and computing the mean Pearson’s correlation across splits. ‘Distance-tuned’ cells were taken as cells that had a ‘split-half correlation’ above 0.5 unless otherwise stated. Across the population of ‘distance-tuned’ cells we observed the majority of cells had a ‘positive’ tuning pattern, firing maximally for a particular distance (reflected by a positive size parameter (*α*) in the gamma fit), many cells however had a ‘negative’ tuning pattern, firing maximally over a range of distances with low firing at a particular distance (reflected by a negative size parameter (*α*) in the gamma fit). We also derived a similar distance-to-goal tuning heatmap for neurons with unique variance explained by distance-to-goal (determined by neGLM model comparisons, see below). When plotting single-unit distance tuning curves a 10 cm Gaussian smoothing kernel was applied. When plotting population distance tuning heatmaps, tuning curves were z-scored, smoothed as above and split into ‘positive’ and ‘negative’ subpopulations. Within each subpopulation, cells were ordered by the maximum (or minimum for ‘negative’ tuning) firing rate of each cell’s fitted gamma-distribution.

### Distance metric comparisons

We made pairwise comparisons between different distance metrics using a Poisson regression framework to determine which distance metric best-explained variance in neural activity. Metrics under comparison were defined as:

- *Manhattan distance to goal*: sum of absolute difference x and y cartesian distances between the subject and the goal, in metres
- *Euclidean distance to goal*: the euclidean distance between the subject and the goal, in metres
- *shortest-path distance to goal*: the geodesic distance between the subject and the goal, that respects maze structure, in metres.
- *future distance to goal*: the distance the subject is yet to travel on their trajectory to goal, in metres
- *path progress to goal*: future distance to goal normalised by the starting distance at cue, a value between 0 and 1
- *time progress to goal*: progress through the total trial duration, a value between 0 and 1

We quantified each metric’s unique predictive power via cross-validated Poisson GLMs, computing a cross-validated coefficient of partial deviance (CPD) for pairwise comparisons of different distance metrics. We first down-sampled navigation time data at 0.5 s resolution, filtering for times in each trial where steps to goal < 30 (off task activity). We then generated a set of 10 gamma basis functions for each distance metric, evenly spaced up to the 90th percentile of observed distances for each metric and trials were then split into five cross-validated folds, within each fold we fit three Poisson GLMs with *L*2 regularisation (determined via nested cross-validation) for each cell: a full model (both metrics), plus two reduced models (each metric alone). On held-out data we computed mean Poisson deviance (*D*) for each cluster, model and fold. Finally computing the CPD for each cell averaged across folds:

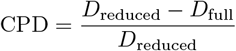

For main pairwise comparisons we took the average CPD over cells for each subject for sessions on Mazes 1 & 2 (where distance metrics were most decorrelated). CPD values from each pair-wise comparison were compared with a paired *t*-test.

### Distance-to-goal decoding

To decode each subject’s shortest-path distance-to-goal from population activity, we first down-sampled and filtered the data the same way as for distance metric comparisons (above), the distances across navigation were partitioned into 5 cm bins uniformly spaced between 0 and 1.6 m and trials were split into 10 cross-validated folds. For each fold we fit a MLR model to predict distance to goal from spike-counts (from both single- and multi-unit activity), under *L*2 regularisation determined from an inner cross-validated fit of the model over the training data, and applied the model to the held out data to return a probability distribution over distance bins for each data sample. For each subject, we performed this decoding on all late sessions and calculated the average predicted probability distribution at each distance to characterise decoding performance. Further we took the maximum decoded probability of this grand-average at each distance bin and compared this to the true distance across subjects.

### Matching neurons across mazes

After spike-sorting sessions individually, units were matched across pairs of sessions recorded in different mazes based on waveform similarity metrics computed using UnitMatch [34]. We used the UnitMatch output to compare distance-to-goal tuning curves of the same cell derived from navigation on different mazes. To quantify if distance-tuning curves across mazes were more similar than chance, we took all pairs of matched distance-tuned cells for each subject and computed the Spearman correlation between their tuning curves. We did the same but over 5000 permuted matches of distance tuned cells, comparing the mean correlation of true matches to the mean correlation of permuted matches across subjects with a paired *t*-test. To illustrate the similarity of distance tuning patterns across mazes, we also constructed distance tuning heatmaps for matched cells. Cells were shuffled between heatmaps to yield heatmap A and heatmap B, containing a mix of cells from Maze 1, Maze 2 and Rooms Maze in each. Cells in heatmap A were ordered as previously described and matched cells in heatmap B followed the same ordering (Figure S8f).

To quantify if place-direction curves across mazes were more similar than chance, similar to the above we took all pairs of matched place-direction tuned cells and compared the true Spearman correlation between matches to permuted controls (correlating only place-directions that were valid on both mazes). To further characterise the low-dimensional structure across matched units we performed a 10-component NMF on population joint place-direction heatmaps across mazes (either for Maze 1 - Maze 2, or Maze 2 - Rooms Maze) and visualised the components as described below (Figure S6e).

### Neural place-direction heatmaps

Neural place-direction heatmaps were calculated by filtering navigation-aligned firing rates for times in the navigation trial phase, where subjects were moving (speed > 0.12 m/s) and were less than 30 steps from the goal, and excluding times at the goal location (immediately before reward). After filtering, firing rates were averaged across frames for each place-direction to give the mean activity across place-directions. For visualisation, markers were placed over each tower or bridge on a maze silhouette where marker colour represents the average firing rate at that location and marker shape represents the directionality of firing at that location. Marker shapes that point in one direction and are flat in all other directions represent complete directional selectivity at that location; markers that point equally in all directions represent equal directional selectivity at that location. Locations with place-directions with less than 0.5 s of occupancy were disregarded and plotted as grey squares. Single neuron place-direction heatmaps were visualised with a custom colour map where low firing rates were coloured to blend into the background maze silhouette and thus accentuate place-directions with higher average firing rates.

Permuted place-direction heatmaps were generated in the same way as above but on firing rates that had been randomly circularly permuted relative to behaviour.

### Neural place-direction representational similarity analysis (RSA)

To quantify the extent to which maze structural features shaped population place-direction representation, we constructed a neural representation dissimilarity matrix (neural RDM) by taking pairwise Pearson’s correlation of population activity vectors across place-direction states. This neural RDM was modelled as the weighted sum of model RDMs designed to capture various maze structural features, including:

- *Euclidean distance*: Pairwise straight-line distances between state coordinates in the 2D maze, to capture whether neural similarity scales with physical proximity in allocentric space.
- *Shortest-path distance*: Pairwise geodesic distances along the maze graph, to capture whether representations reflect maze, rather than purely Euclidean, proximity.
- *Boundary distance*: Absolute differences in each state’s distance to the nearest wall. Captures sensitivity to proximity to environmental boundaries.
- *Betweenness centrality*: Absolute differences in graph-theoretic betweenness centrality between states, to capture neural tuning to central routes and dead-ends.
- *Decision point distance*: Absolute differences in shortest-path distance to the nearest decision point (degree 3 or 4 node), to capture organisation relative to behaviourally relevant choice point/bottleneck locations.
- *Corner identity*: Binary dissimilarity (0 = same corner, 1 = different corners), to capture discrete tuning to the four corners of the maze.

For each subject we fit a multiple linear regression model predicting neural dissimilarity from the vectorised RDMs above, where the shortest-path regressor was orthogonalised relative to the Euclidean regressor as these features were fairly collinear. Subject betas for each RDM were compared to zero using a one-sample *t*-test (with BH correction).

### Dimensionality reduction on population place-direction tuning

To quantify the low dimensional structure in population place-direction tuning, we constructed a data matrix of n neurons by n place-directions (*X*_*N*_) and performed principal component analysis (PCA) or non-negative matrix factorisation (NMF) to yield a set of neural components (*K*_*N*_) and neural loadings (*L*_*N*_):

**PCA:**

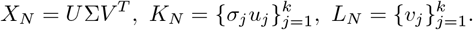

**NMF:**

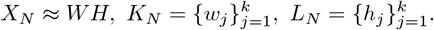

Before NMF or PCA, missing values in the neural data matrix (unvisited place-directions) were replaced with the mean across valid place-directions and all heatmaps were normalised to length one. NMF with 10 components was performed for visualisation purposes, whereas PCA was used for various quantitative analyses.

To confirm that the dimensionality of neural place-direction tuning was lower than chance (expected by pure autocorrelation in neurons and behaviour), we compared the cross-validated cumulative variance explained by the first *k* principal components of real place-direction tuning with cumulative variance explained by the first *k* principal components of permuted place-direction heatmaps. Variation across subjects was estimated with 1000 bootstrapped resamples of subjects whose real or permuted data went into the variance explained analysis. To quantify the difference in dimensionality between conditions we compared the area under cumulative variance explained curves across permutations (higher area = lower dimensional).

### Dimensionality reduction on behavioural trajectories

Trial-by-trial behavioural trajectories could be represented in the same place-direction space as neural heatmaps. Here, navigation trials were represented as a binary vector corresponding to the place-directions visited on a given navigational trial (filtering for only the last 30 steps to goal, same as place-direction heatmaps). These vectors were concatenated to build a behavioural data matrix of n trials by n place-directions, place-direction values in each trial were normalised to length one and PCA or NMF was performed (as above) to generate a set of behavioural components and behavioural loadings. NMF with 10 components was performed for visualisation comparison with the neural NMF, whereas PCA was used for various quantitative analyses.

### Relating low-dimensional structure of neural place-direction tuning and subjects’ behaviour

To test whether mFC place-direction tuning reflected the low-dimensional structure of behaviour— and thus formed an efficient code for behavioural trajectories—we asked whether the principal components of behaviour also explained variance in neural place-direction tuning. We performed PCA on the behavioural matrix (*X*_*B*_), yielding orthonormal behavioural components (the columns of *V*), and projected the neural matrix (*X*_*N*_) onto the first *k* of these to compute the cumulative fraction of neural variance explained, CVE(*k*):

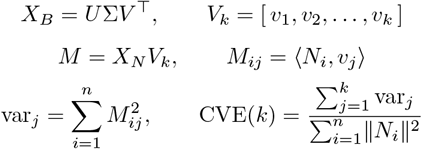

where *N*_*i*_ is the (mean-centred) place-direction tuning of neuron *i* and *n* the number of neurons.

We computed CVE(*k*) not only for the subjects’ true behaviour but also for synthetic behaviour, matched trial-by-trial to the real data (same start and goal locations) but with decisions drawn from either a random-walk or an optimal policy. Variance explained was estimated throughout with split-half cross-validation.

Repeating this across all subjects gave a mean CVE curve for each condition. To estimate cross-subject variability, we bootstrap-resampled subjects (with replacement; 1000 resamples) and recomputed each curve. For every resample we measured the area under the curve (AUC), giving a bootstrap AUC distribution per condition (neural data, true, random, and optimal behaviour). Two conditions were taken to differ significantly when their AUC distributions overlapped by less than 5% (*p <* 0.05).

### Future/past place-direction decoding

We asked whether mFC population activity encoded a subject’s past and future behaviour on individual trials (occupied place-directions). For this analysis we focused on timepoints during navigation where subjects were approaching (for future decoding) or leaving (for past decoding) decision points on the maze (nodes of degree 3 or 4; Figure S5h). From 0.1 s binned navigation aligned spike-counts, we encoded either just subject’s current place-direction (coded as a one-hot vector) or both subject’s current place-direction and mFC spike-counts and performed cross-validated multinomial logistic regression to predict subject’s place-direction one, two, three, etc. steps in the past/future, where 1 step was equivalent to a tower-tower transition. We then calculated the additional decoding accuracy afforded by the inclusion of mFC spike-counts at each timestep by comparing decoding conditions. Above chance decoding was determined by comparing the subject average decoding accuracy between control and spike-count decoders with BH-corrected paired *t*-tests across timepoints.

### neGLM model architecture

To quantify how neural activity encodes behavioural variables like location and distance, we developed a new neural network-based embedding model that combines data across sessions. This is necessary because the same neurons are not recorded across all sessions, but a single session contains too little data to regress neural activity onto high-dimensional behaviour. Instead, we project the behaviour into a learned lower-dimensional *latent space* that is shared across *all* sessions. The only session-specific parameters are linear mappings that specify how the activity of neurons recorded in that session depends on the latent variables.

More specifically, we used a ReLU neural network to generate latent variables 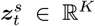 at time *t* in session *s* from an input vector 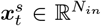 that indicates the observations at time *t* in session *s*:

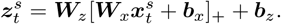

Here, 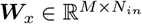 is the mapping from inputs to the hidden layer, and ***W***_*z*_ ∈ ℝ^*K*×*M*^ maps from the hidden layer to the latents. ***b*** are the corresponding bias vectors. Importantly, these parameters are all shared across sessions. We used a linear exponential Poisson model to model the dependency of neural activity on the latents,

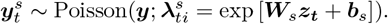

Here, 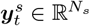 is the binned neural activity at time *t* in session *s*, and ***W***_*s*_ and ***b***_*s*_ are session-specific parameters that indicate how neural activity depends on the latents. The model parameters were optimised by minimising the negative log likelihood of the data:

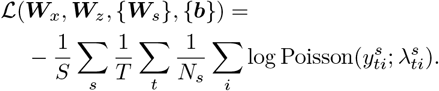

Here, subscript *i* indexes a single one of the *N*_*s*_ neurons recorded in session *s*. All parameters were optimised using Adam with a learning rate of 1 × 10^−2^ and a weight and activity regularisation strength of 1 × 10^−1^ on the embedding network, across 3000 epochs of the dataset. Model performance was stable across a range of hyperparameter values, and training-loss and train/test performance curves indicate that networks converged (Figure S9f-h).

To study the interactions between two sets of input variables ***x***_1_ and ***x***_2_, we compared two different models that either combined or partitioned the input streams. In the non-partitioned ‘nonlinear interaction’ model, ***x*** = [***x***_1_, ***x***_2_] is the concatenation of the input variables. In the partitioned ‘linear interaction’ model, the input variables are embedded separately. The latents ***z*** = ***z***_1_ + ***z***_2_ are then computed as the sum of latents arising from separate sets of embedding parameters applied to the separate input channels (Figure S9a). This linear model only allows for linear interactions in the latent space, corresponding to multiplicative interactions in the firing rates after the Poisson nonlinearity. In particular, the predicted firing rates in the nonlinear model are given by

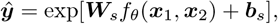

for some learned function *f*_*θ*_. In the linear model, the predicted firing rates are given by

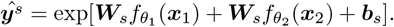

If these two models explain the data equally well, we conclude that the corresponding input variables interact linearly in latent space. This would for example preclude conjunctive tuning for ***x***_1_ and ***x***_2_.

To quantify the fit of the embedding models, we used a nested cross-validation with an outer split across sessions and an inner split across trials. In the outer loop, we held out one session at a time, fitting the shared embedding parameters {***W***_*x*_, ***W***_*z*_, ***b***_*x*_, ***b***_*z*_*}* on the remaining *S* − 1 sessions and applying them to embed the behaviour of the held-out session. Because this session’s neurons were never seen during embedding, its session-specific parameters {***W***_*s*_, ***b***_*s*_*}* had to be estimated from its own data. In the inner loop, we fitted these on 80% of the held-out session’s trials and computed the deviance explained, across all of its neurons, on the remaining 20%. Rotating the inner split over every trial fold and the outer hold-out over every session, we averaged the deviance explained over all session-by-fold combinations.

### neGLM model comparisons

We compared performance across models with paired, BH-corrected *t*-tests across subjects, computed on each subject’s cross-validated deviance explained, averaged across all of that subject’s neurons (pooled over sessions and mazes). We computed CPD as described above, but from the neGLM’s cross-validated deviance scores. We classified a neuron as encoding a given variable when its per-fold CPD values, computed across 10 cross-validation folds, were significantly greater than zero (onesided one-sample *t*-test, *p <* 0.01). We did not correct for multiple comparisons, so the expected per-comparison false-positive rate is approximately 1%. For our final set of neGLM comparisons, we progressively enlarged the input set until the explainable variance saturated, introducing a new variable only when it could not already be computed from nonlinear interactions among existing inputs. This restriction is necessary because of the flexibility of the neGLM embedding. For example, given enough training data, a model supplied with place and goal could compute distance-to-goal within the embedding and explain variance through it, even when no explicit allocentric goal representation is present. Restricting each addition to variables not already derivable from existing features ensured that every new input probed for genuinely new explanatory structure rather than a quantity the embedding could already construct.

### neGLM mixed selectivity permutation test

To test whether the unique variance attributed to each input stream ***x***_1_ and ***x***_2_ was carried by distinct neuronal subpopulations (factorised coding) or distributed across neurons (mixed coding), we constructed a null distribution under random rotations of the held-out population activity. For each heldout session *s*, we collected the binned spike data into a matrix 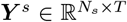 with columns 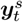, and rotated across neurons by a Haar-uniform random orthogonal matrix ***Q*** ∈ *O*(*N*_*s*_):

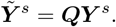

***Q*** was obtained from the QR decomposition of a matrix of i.i.d. standard normals after correcting the signs of the diagonal of ***R*** to ensure uniform sampling on the orthogonal group [97]. This rotation preserves the population-level covariance of ***Y*** ^*s*^ while scrambling each neuron’s alignment to the variable axes.

Because 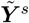 is real-valued and can take negative entries, the exponential-Poisson readout in ***W***_*s*_, ***b***_*s*_ is no longer well-defined. We therefore replaced the session-specific readout with a Gaussian-likelihood ridge regression from the latents 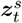 to 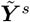, fit on 80% of trials in session *s*, and reported *R*^2^ on the remaining 20%. For an apples-to-apples reference, an identical ridge readout was also fit on the unrotated spikes ***Y*** ^*s*^ (Figure S9b).

For each held-out fold we computed the perneuron unique variance explained 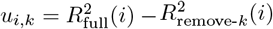. for input stream *k* ∈ {1, 2}.Each permutation was then summarised by a per-subject axis-alignment score,

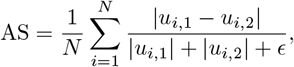

averaged across subjects. Repeating with 1,000 rotations per fold yielded a null distribution over the subject-mean axis-alignment score, against which two-sided *p*-values were computed (Figure S9b).

### Egocentric action tuning

Prompted by the unique variance explained by egocentric actions in the neGLM model comparison, we sought to unpack egocentric-action tuning in the data. We derived tuning curves by aligning each neuron’s firing rate to the moment a subject entered a tower to make an action, split by action type: turn left, turn right or go forward (we ignored go-back actions, which occurred rarely). We plotted single-neuron tuning as the mean firing rate across actions, smoothed with a 0.2 s Gaussian kernel, and computed a split-half correlation metric as described for the distance-to-goal and place-direction tuning curves. For the population heatmaps, we grouped all egocentric-action-tuned neurons (split-half correlation *>* 0.3) into three blocks by preferred action, then ordered neurons within each block by their time of maximum (non-cross-validated) tuning (Figure S10a&b).

To visualise the population dynamics aligned to egocentric actions, we computed each unit’s average activity for each action and concatenated these across actions into a data matrix of shape [*n*_neurons_ × (*n*_actions_ × *n*_timepoints_)]. We performed PCA on this matrix to identify the subspace capturing the most variance across action types, then projected each action’s average population activity into this subspace to visualise neural trajectories, further splitting each action into free and forced variants (Figure S10c). Free actions occurred at towers with three or four connected bridges, where subjects chose how to proceed; forced actions occurred at towers with one or two bridges, where they could only continue forward or turn around.

### Velocity tuning

Movement velocity was also shown to explain unique variance in mFC from neGLM model comparisons. To characterise this tuning further we calculated velocity tuning heatmaps by calculating average firing rate over binned instantaneous velocities (*v*_*x*_, *v*_*y*_) during navigation for each cell. Similar to other tuning curves, we derived a split-half correlation metric to find cells with stable velocity tuning (> 0.65; ‘velocity tuned cells’). The tuning of these cells ranged from a single firing field to multiple, symmetrically arranged fields. To quantify this structure across the population we calculated a rotational autocorrelogram for each velocity tuned cell:

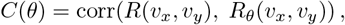

where *R*_*θ*_(*v*_*x*_, *v*_*y*_) denotes the velocity tuning map rotated by angle *θ*. We further decomposed these autocorrelograms into a power spectrum of rotational symmetries using a discrete Fourier transform:

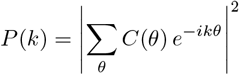

where *k* ∈ {1, 2, 3, 4} indexes harmonic order. These harmonic profiles were used to visualise symmetrical structured tuning across the population, ordering neurons by their dominant harmonic (Figure S11).

### Egocentric angle to goal tuning

To characterise single-neuron tuning to the egocentric angle to goal — the angle between the animal’s current heading and the vector pointing to the goal location (0^◦^ = goal ahead, 90^◦^ = left, 180^◦^ = behind, 270^◦^ = right) — we binned this variable into 120 equally-spaced bins over [0, 360)^◦^ and, for each cluster, computed the mean firing rate per trial × per bin during navigation. Single-neuron tuning curves were obtained by averaging across trials within each bin with an 8^◦^ circular smoothing kernel applied for visualisation. Cells with stable tuning were identified by another split-half correlation metric; we retained ‘egocentric-angle-to-goal tuned’ cells with a split-half correlation *>* 0.6 for further analysis (Figure S12a).

To summarise tuning across the population, we fit a four-parameter von Mises function to each neuron’s tuning curve,

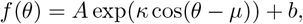

where *µ* is the preferred angle, *κ* the concentration, *A* the amplitude and *b* a baseline offset. Fits were cross-validated by fitting on one trial-half and evaluating *R*^2^ on the held-out half across the same 50 splits used for the split-half correlation metric, with the per-neuron *µ* reported as the circular median across splits. As a non-parametric measure of tuning concentration that does not depend on the parametric fit, we additionally computed the mean resultant length (MRL) of the smoothed tuning curve (0 = uniform, 1 = perfectly concentrated). Population heatmaps and circular histograms were organised by cross-validated preferred angle *µ*, and the joint distribution of *µ* and MRL was used to summarise the population tuning structure (Figure S12b–d).

### Population theta-modulation

To quantify theta-modulation across the population we extracted theta-power from the LFP signal (bandpass filter: 7-11 Hz), and counted the number of spikes in each of 12 binned theta-phases (Figure S13f), during periods of moving navigation. These modulation profiles were normalised to the mean across theta bins and averaged over all neurons, or over specific subpopulations with unique variance explained by place-direction or distance-to-goal (from neGLM model comparisons; Figure S13c). Theta modulation was quantified by fitting a sinusoid to each subject’s modulation profile and then submitting the resultant amplitude and phase parameters to a Hotelling’s *T* ^2^ test. Phase-offsets across population modulation curves were quantified by calculating per-subject offset differences and testing those values for non-uniformity with Rayleigh’s test.

### Decision-aligned theta-power

To ask whether mFC theta power was stronger before decisions where subjects needed to use knowledge of maze structure over habitual strategies (shown to be mFC dependent in the optogenetic inhibition experiment), we aligned theta-band power to decision points where structure-based and habitual strategies were in conflict (structure ≠ habit) stratified by whether subjects chose structure-based (optimal) or habitual (error) actions. We compared differences in theta power pre-decision across subjects with paired *t*-tests at successive pre-decision timepoints (BH-corrected). Because structure-based and habitual choices at these decision points likely differed in average speed and distance-to-goal, we confirmed differences using a per-timepoint linear regression of theta power on speed, distance-to-goal, and a choice (habit vs. structure) regressor (+1 chose-structure, −1 chose-habit), testing the per-subject coefficients against zero across the group (*t*-tests *>* 0 speed, *>* 0 choice, two-sided distance), either within a hypothesised window or over successive timepoints aligned to the decision.

### Theta-modulated distance-to-goal decoding

To test for theta modulation in the distance-to-goal signal we used a similar MLR decoding framework as previously described but where population activity (spikes) had been stratified by theta phases and then down-sampled to 0.5 s resolution (Figure S13f), yielding 12 neuron × timepoint population activity matrices and one additional reference average activity matrix taken as the mean spike count per neuron per timepoint over theta phases (Figure S13d). In each cross-validated fold, the model was trained to predict distance-to-goal from reference average data and tested on held out data from the spike-counts from each theta-phase bin separately, taking the decoded distance as the sum of distance-bin centres weighted by their predicted probabilities. We performed this decoding on all sessions for each subject on data less than 0.8 m from the goal where distance decoding was most accurate. For each subject, we then calculated the decoding error as the true minus decoded distance at each sample and theta-phase and took the average of this decoding error across samples for each theta-phase to yield distance-to-goal decoding bias curves for each subject. Decoding bias curves were then demeaned before further averaging across subjects and statistically compared as described above (Hotelling’s *T* ^2^ test).

### Theta-modulated distance-to-goal tuning

To test for theta modulation outside the decoding framework, we computed single-unit distance-to-goal tuning curves (4 cm bin size, as described above) separately for spikes assigned to each of 12 theta-phase bins, yielding 12 phase-specific tuning curves per neuron. A reference tuning curve was also computed by averaging spikes across all theta phases. Each neuron’s tuning curves (phase-specific and reference) were normalised by the maximum firing rate of its reference curve, then upsampled to 0.5 cm resolution by quadratic interpolation. For each subject and theta-phase bin, the tuning curves of all neurons with strong distance-to-goal tuning (split-half correlation *>* 0.7) were stacked to form a theta-phase-specific neural tuning matrix (neurons × distance bins), with a corresponding reference tuning matrix constructed from the same neurons’ reference curves. For each phase, we computed the mean squared error between the theta-phase-specific and reference tuning matrices across a range of relative x-shifts (−8 to +8 cm), and took the shift minimising MSE as the optimal alignment. The resulting set of 12 optimal shifts defined a theta-modulation profile for each subject, which was tested for significant modulation using a Hotelling’s *T* ^2^ test (as described above).

### Theta-modulated place-on-trajectory decoding

To test whether the place-direction representation was theta-modulated we used an approach analogous to the theta-modulated distance-to-goal decoding described above, but with current maze location (place) as the decoded variable. Population spike counts were summed over a 0.4 s window and sampled every 0.1 s. Cross-validated place-decoder outputs at each theta-phase (a probability distribution over maze locations) were transformed into a trajectory-projected place bias. For each sample *t* and theta-phase *ϕ* we restricted the place-decoder probabilities 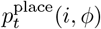 to the animal’s current location and the two locations immediately ahead and behind along its trajectory (*L*_*t*_), renormalised them to sum to 1 within *L*_*t*_, and weighted by the signed trajectory distance *d*_*t*_(*i*) from the current location:

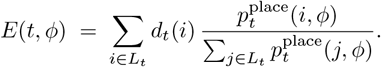

Trajectory-projected place biases were then averaged across samples within each theta-phase to yield a place-on-trajectory decoding bias curve for each subject. Bias curves were demeaned before averaging across subjects as described above. Phase offsets between the decoded distance-to-goal and decoded place-on-trajectory modulation profiles were computed for each subject as the circular difference between the trough phases of sinusoids fitted to the two profiles. The distribution of phase offsets across subjects was tested for non-uniformity using Rayleigh’s test.

### Behavioural-timescale correlation of place and distance-to-goal decoding errors

To test whether place and distance-to-goal representations fluctuated together on a behavioural timescale, we trained two independent MLR decoders on non-overlapping populations of selectively tuned units (place-direction or distance-to-goal tuned, defined via the neGLM variance-explained criteria described above): a place decoder predicting the animal’s current maze location and a distance-to-goal decoder predicting its distance to goal. Population spike counts were sampled at 0.2 s and decoding was performed using leave-one-trial-out cross-validation. For each held-out sample (*t*) we computed, from each decoder, a trajectory-projected bias *E*_*t*_ — the signed-distance-weighted sum of decoder probabilities over *L*_*t*_ as defined above without stratification by theta phase (for the distance decoder, evaluated over the corresponding step-distances to goal) — yielding place and distance-to-goal bias time courses 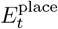 and 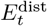. We then computed, per subject, the Pearson correlation between 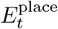 and 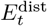 across samples from late sessions (not including data from sessions whose place or distance-to-goal decoders failed to clear twice chance accuracy). Subject-level correlations were tested against zero with a one-sided one-sample *t*-test.

## Supplementary Figures

**Figure S1:**
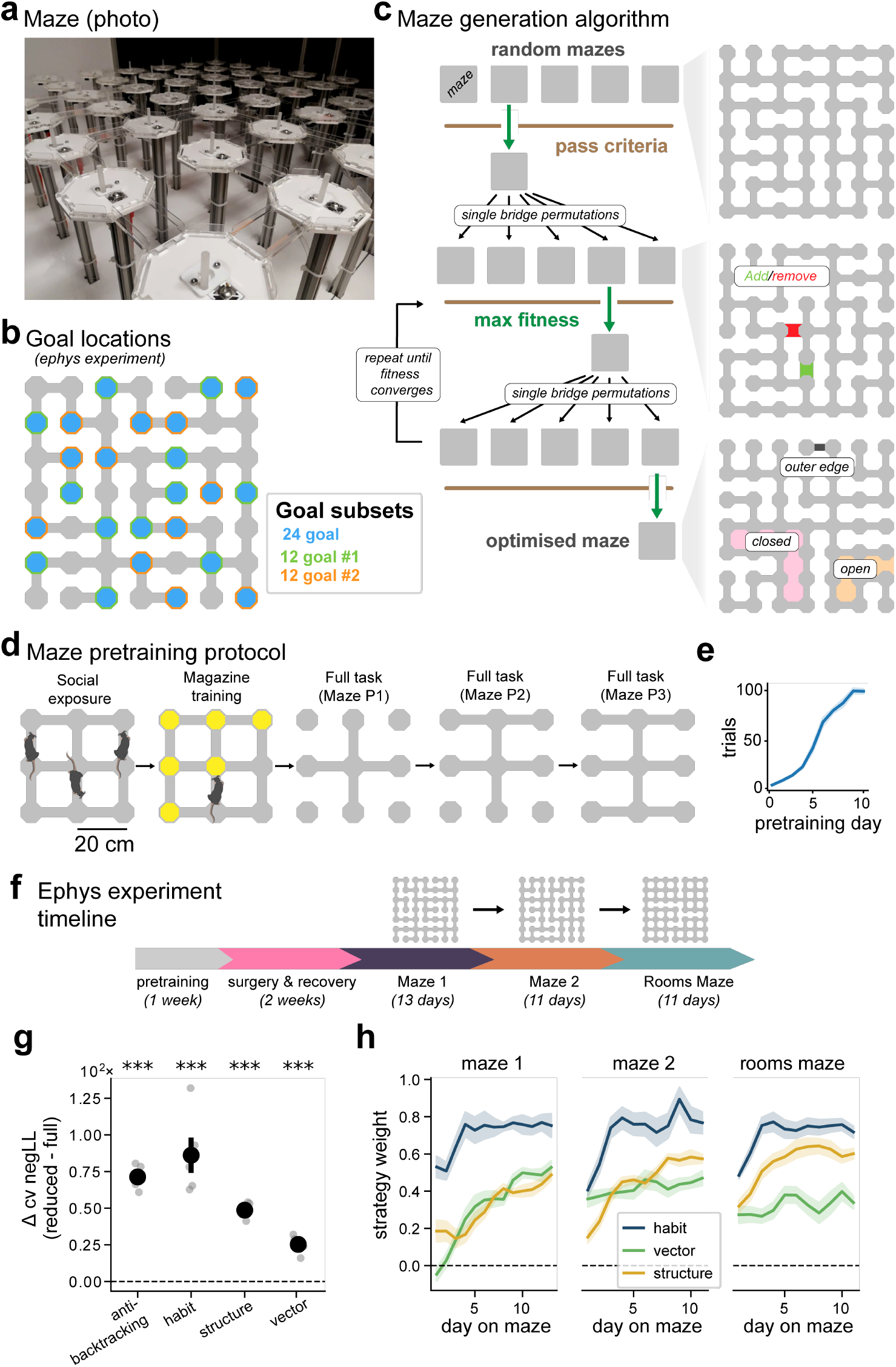
Maze Design and Behaviour. **(a)** Elevated grid maze apparatus (photo). Note mice perform the task in the dark. **(b)** Goal subset locations used for the electrophysiology experiment. Note the same goal locations and goal subsets were consistent across maze configurations. **(c)** Left: schematic of maze generation algorithm. Right: Example random, mutated & optimised mazes (top to bottom). **(d)** Pre-training protocol on 3-by-3 maze. **(e)** Trials per session over pretraining days. **(f)** Visualisation of the ephys experiment timeline. After pretraining and surgical implantation of electrodes, mice learned to navigate three complex mazes in sequence, from naive to expert on each maze. **(g)** Cross-validated negative log-likelihood (cv negLL) estimates from full and reduced model (all but one strategy) comparisons demonstrate that each navigation strategy contributes uniquely to model fit, justifying the inclusion of all strategies. One-sample *t*-tests *>* 0 (***p < .001). **(h)** Habitual choice (habit), vector-based (vector), and structure-based (structure) navigation strategy weights from the mixture of strategies model fit to subjects’ navigational decisions separately for each maze-day. Maze 1 (left), Maze 2 (middle) and Rooms Maze (right). Error bars and shading represent sem across subjects.

**Figure S2:**
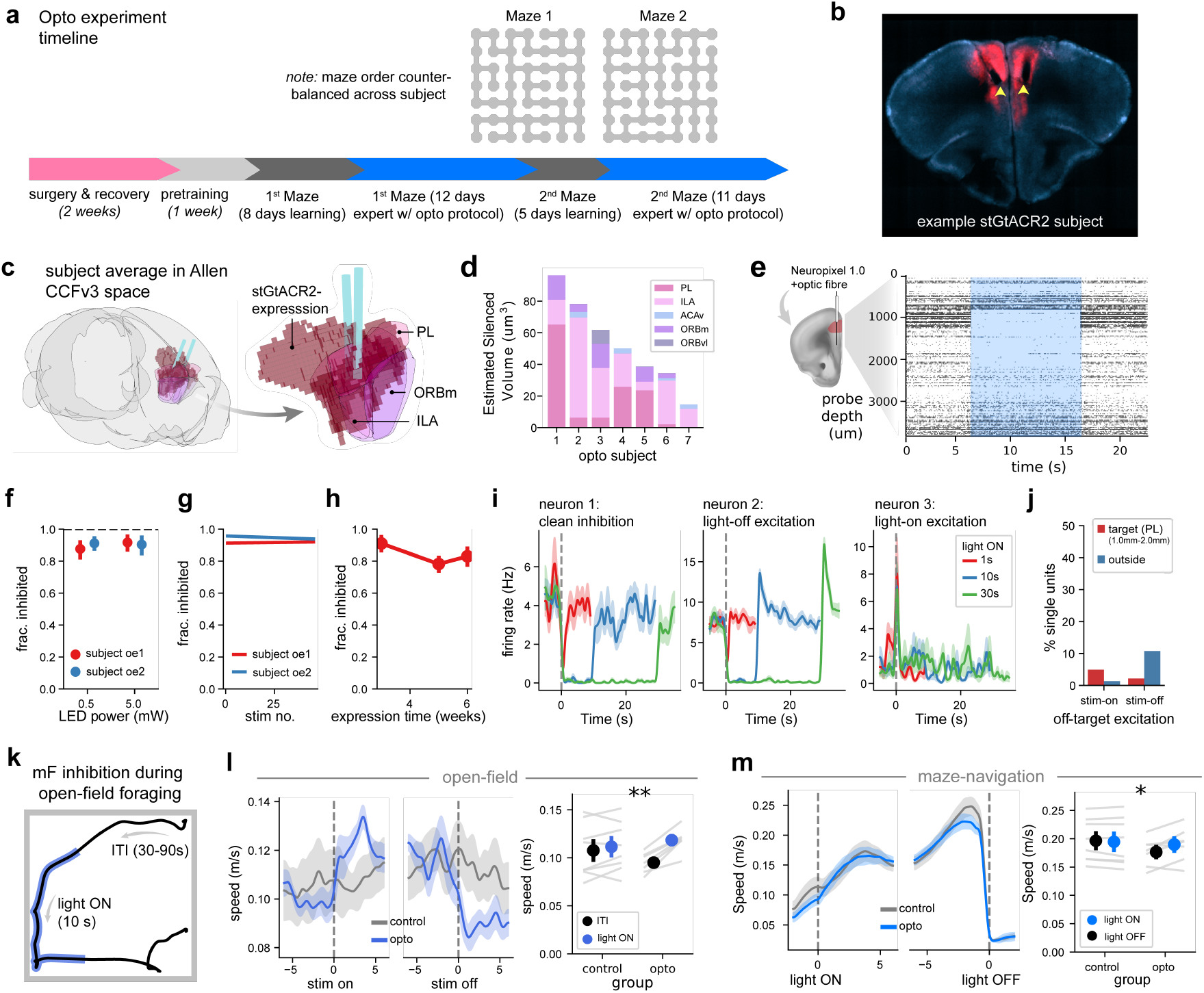
Optogenetic silencing of mFC. **(a)** Visualisation of the optogenetic experiment timeline. After surgery and pretraining, mice were split into groups that experienced Maze 1 then Maze 2 or Maze 2 then Maze 1 (counterbalanced). **(b)** Example histological section (blue) illustrating stGtACR2 expression (red) and optic-fibre placement (tips marked with yellow arrows) in mFC. **(c)** Brains were imaged with serial 2p tomography and registered to the Allen mouse atlas using brain-globe. Viral expression was averaged across subjects, thresholded, denoised, and plotted with the average optic-fibre placement and anatomical reference regions (PL: prelimbic, ILA: infralimbic, ORBm: medial orbito-frontal) in Allen CCFv3 coordinates (brain-render). **(d)** mFC inhibition occurred primarily in the prelimbic and infralimbic subregions. The total volume, and corresponding anatomical regions, of mFC inhibition in each subject were estimated as the union of two cones extending from fibre tips (length: 500 µm, half-angle: 35°) and thresholded viral expression in atlas space. **(e)** Two additional subjects received the stGtACR2 construct and a custom implant housing a Neuropixel 1.0 probe and optic fibre in mFC. Three weeks after surgery, electrophysiological recordings with simultaneous blue-light (473 nm) delivery to mFC (1, 10 or 30 s pulses, 45–90 s inter-trial interval) were conducted in the subjects’ home cages over 3 weeks, to assess the efficacy of our optogenetic approach. **(f)** We quantified the fraction of mFC neurons inhibited in our target region (1000–2000 µm from the brain surface roughly corresponding to the prelimbic subregion). We found that both 0.5 mW and 5 mW laser powers inhibited the majority of target cells. **(g)** Inhibition stayed relatively stable over 30 min of repeated light-ON events within a session. **(h)** Inhibition remained efficacious over the three-week recording period. **(i-j)** While the majority of targeted cells exhibited clean inhibition during light-ON (h left), some cells, mostly in the target region (i red bars) experienced rebound excitation as the light was turned off (h middle) and some cells, mostly outside the target region (i blue bars), were transiently excited as the light was turned on (h right). **(k)** We also performed mFC inhibition (in main cohort subjects) during open-field foraging, where mice explored an open-field to find randomly scattered chocolate sprinkles over 30 min sessions with 10 s light-ON periods separated by 30–90 s inter-trial interval periods. **(l)** We observed a subtle and transient increase in the subject’s movement speed aligned to stim in opsin-expressing animals (left) which translated to a significant increase in average speed during light-ON compared to controls (right; Mixed ANOVA group ×light-ON: *F* (1, 13) = 15.96, *p* = .0015, 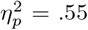). **(m)** A similar subtle, but significant, increase in movement speed during light-ON was observed in the opto-group during maze navigation (right; Mixed ANOVA group×light-ON:*F* (1, 13) = 7.58, *p* = .0164 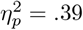).

**Figure S3:**
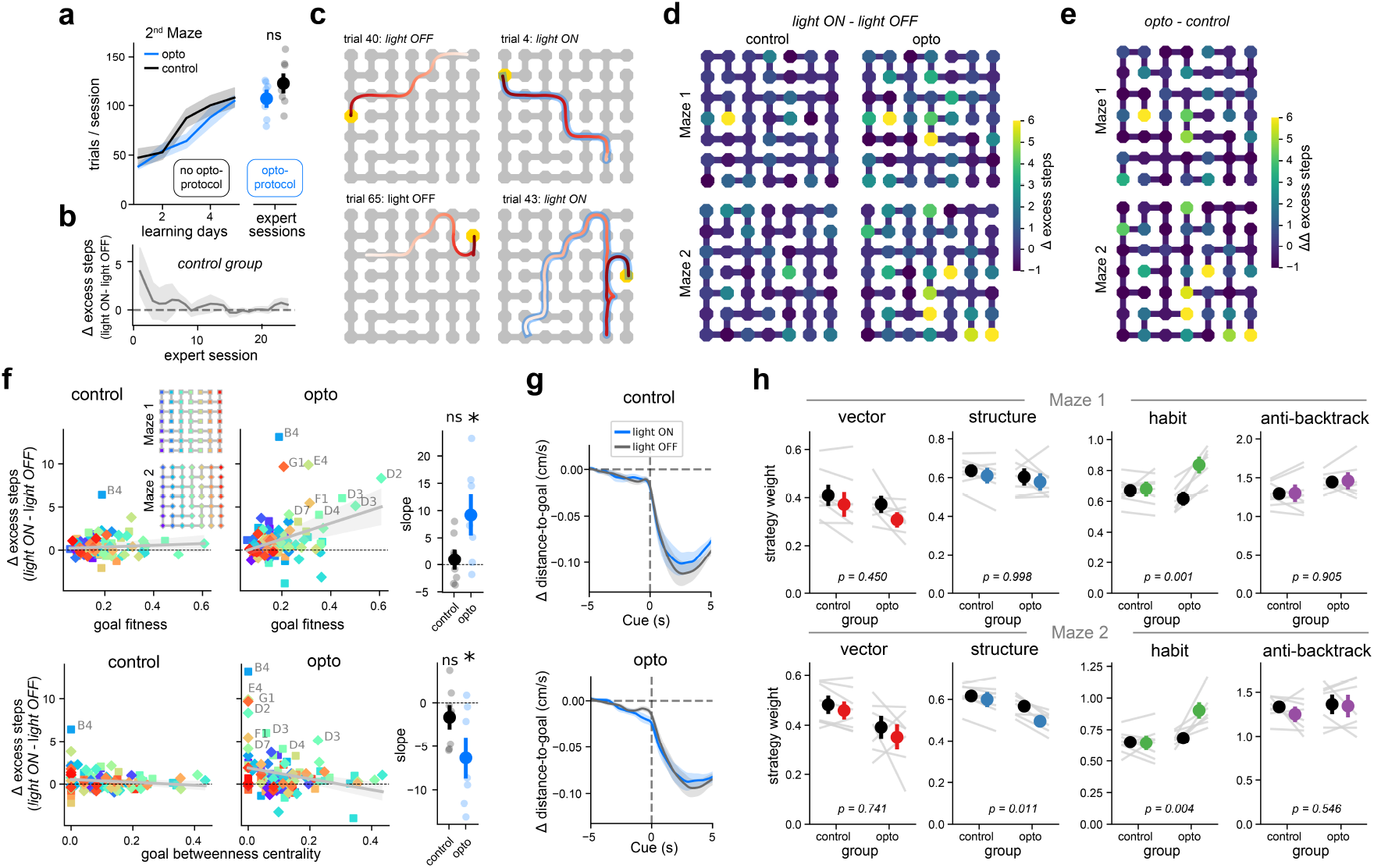
Silencing mFC during maze navigation. **(a)** Trials per session increased over learning (left: linear mixed model: day: *β* = 8.4, *z* = 16.6, *p <* .001, maze×day: *β* = −2.1, *z* = −8.2, *p <* .001) and remained stable over expert behaviour (right: t-test across groups: *T* (13) = 1.41, *p* = 0.182) on the 2nd maze. Note maze order was counterbalanced across subjects, so that some subjects experienced Maze 1 then Maze 2 and others Maze 2 then Maze 1. Formatted as in Figure 2c. **(b)** Subjects adapted to light-ON trials over 5 days, so early expert sessions were excluded from summary analyses. Plotted: difference between session average excess steps in light-ON and light-OFF trials over all days of expert behaviour (opto protocol active). **(c)** More example navigation trajectories in an opsin-expression subject during light-OFF (left column) and light-ON (right column) trials. On some light-ON trials subjects navigated to goals efficiently (e.g. right top), but on others, where they needed to deviate from the maze’s central routes, paths to goal were much less efficient (e.g. right bottom). Formatted as in Figure 2g. **(d)** mFC inhibition impaired navigation to some goals more than others. Plotted: difference in average excess steps between light-ON and light-OFF per goal (edges plotted in dark blue; value = 0). Heatmaps represent the average across subjects in the control (left) and opto (right) groups on Maze 1 (top) and Maze 2 (bottom). **(e)** Difference between opto and control heatmaps in panel d, to visualise which goals accrued more excess steps in light-ON trials relative to the controls. Formatted as in panel d. **(f)** This pattern of excess steps across goals was systematically related to measures of goal fitness (the decorrelation of Euclidean and shortest-path distances between all pairs of points on the maze including the goal) and goal betweenness centrality (the sum of the fraction of all-pairs shortest-paths that pass through the goal). Plotted: Difference in excess steps between light-ON and light-OFF trials per goal, further stratified by goal-fitness (top row) or goal betweenness centrality (bottom row) for control (left) or opto (middle) subjects. Points represent cross subject averages and are coloured by maze legend in the top-right inset (Maze 1: square marker, Maze 2: diamond marker). Grey fitted line and shaded error represents mean ± sem across subjects of linear regression fits to the data. Slopes from these were significantly greater than zero for goal-fitness (top-right; one-sample *t*-test per group. control: *T* (7) = 0.63, *p* = 0.550, opto: *T* (6) = 2.70, *p* = 0.035), and significantly less than zero for goal betweenness centrality (bottom-right; one-sample *t*-test per group. control: *T* (7) = −1.41, *p* = 0.200, opto: *T* (6) = −2.97, *p* = 0.025), in opsin-expressing subjects but not controls. **(g)** Opto and control groups were similarly goal-directed at the onset of navigation across light-ON and light-OFF trials, suggesting that mFC inhibition did not lead to a general increase in task disengagement. Plotted: rate of change of distance to goal aligned to cue, plotted separately for control (top) and opto (bottom) groups. **(h)** Mixture of strategies modelling with vector-based, habit, anti-backtracking and structure-based strategies was fit to subjects’ navigation decisions on light-ON and light-OFF trials. Group × light-ON interaction p-values (Mixed ANOVA) printed at the bottom of each subpanel. Plots formatted as in Figure 2d, coloured markers indicate average weights fit to light-ON trials.

**Figure S4:**
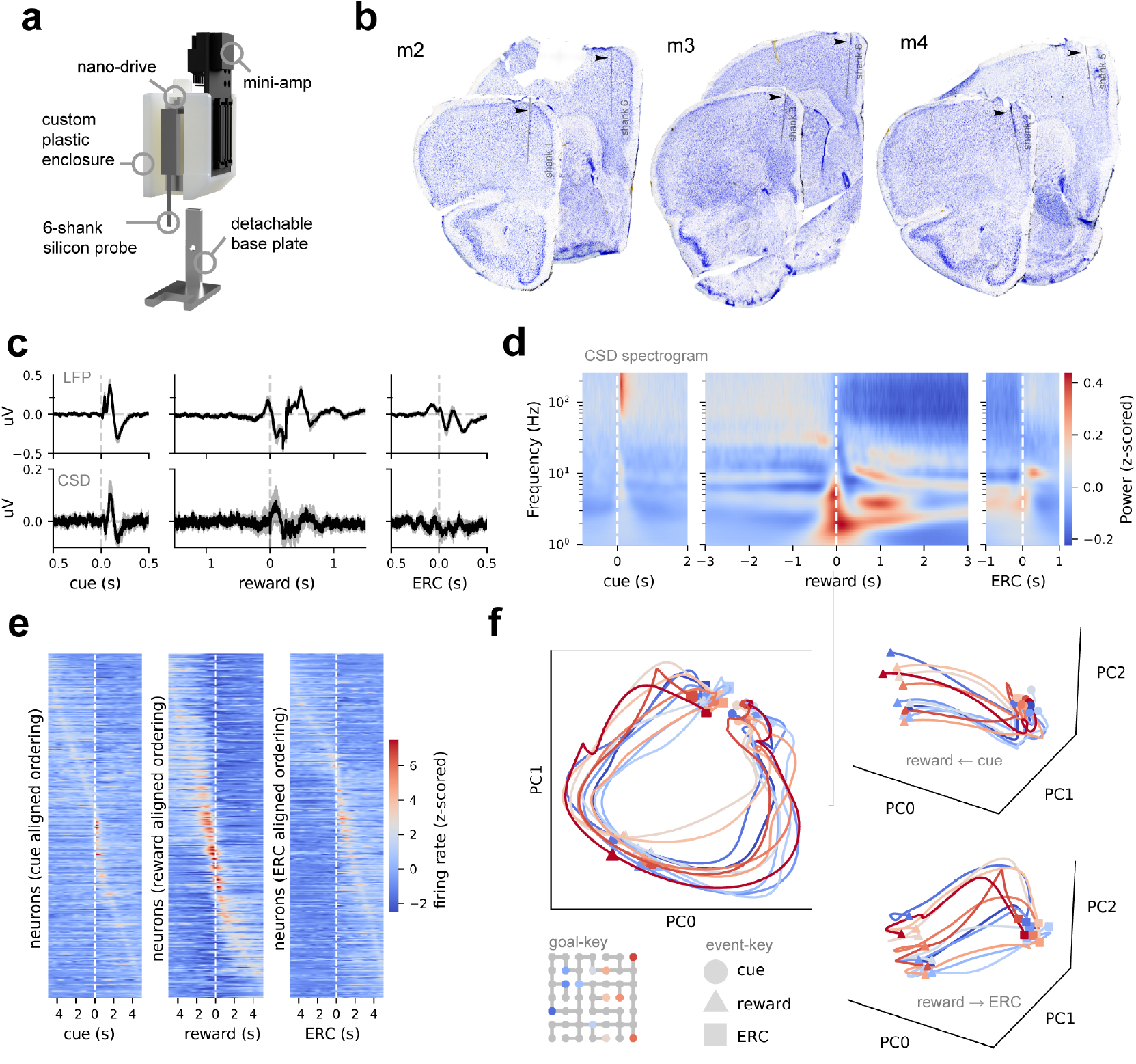
Neural representations of task structure. **(a)** Custom silicon probe implant with labelled components. **(b)** Example histology images from three subjects (with example anterior and posterior shanks), probe tracts are marked in light grey and a black arrow. **(c)** Average LFP (top row) and CSD (bottom row) signal aligned to cue (left), reward (middle) and end-reward-consumption (ERC, right), shaded area represents standard error across subjects. **(d)** Mean power spectral density of CSD signal (z-score normalised across frequencies) aligned to cue (left), reward (middle) and end-reward-consumption (ERC, right). **(e)** Population activity heatmaps aligned to cue (left), reward (middle) and end-reward-consumption (right). Note that single units are ordered independently in each heatmap. **(f)** Population activity projected onto the first three principal components of goal-stratified, trial-averaged single-unit firing rates, with each trajectory coloured by navigational goal. PC1 and PC2 capture rotational dynamics in which trial progression (cue → reward → ERC) traces a loop.

**Figure S5:**
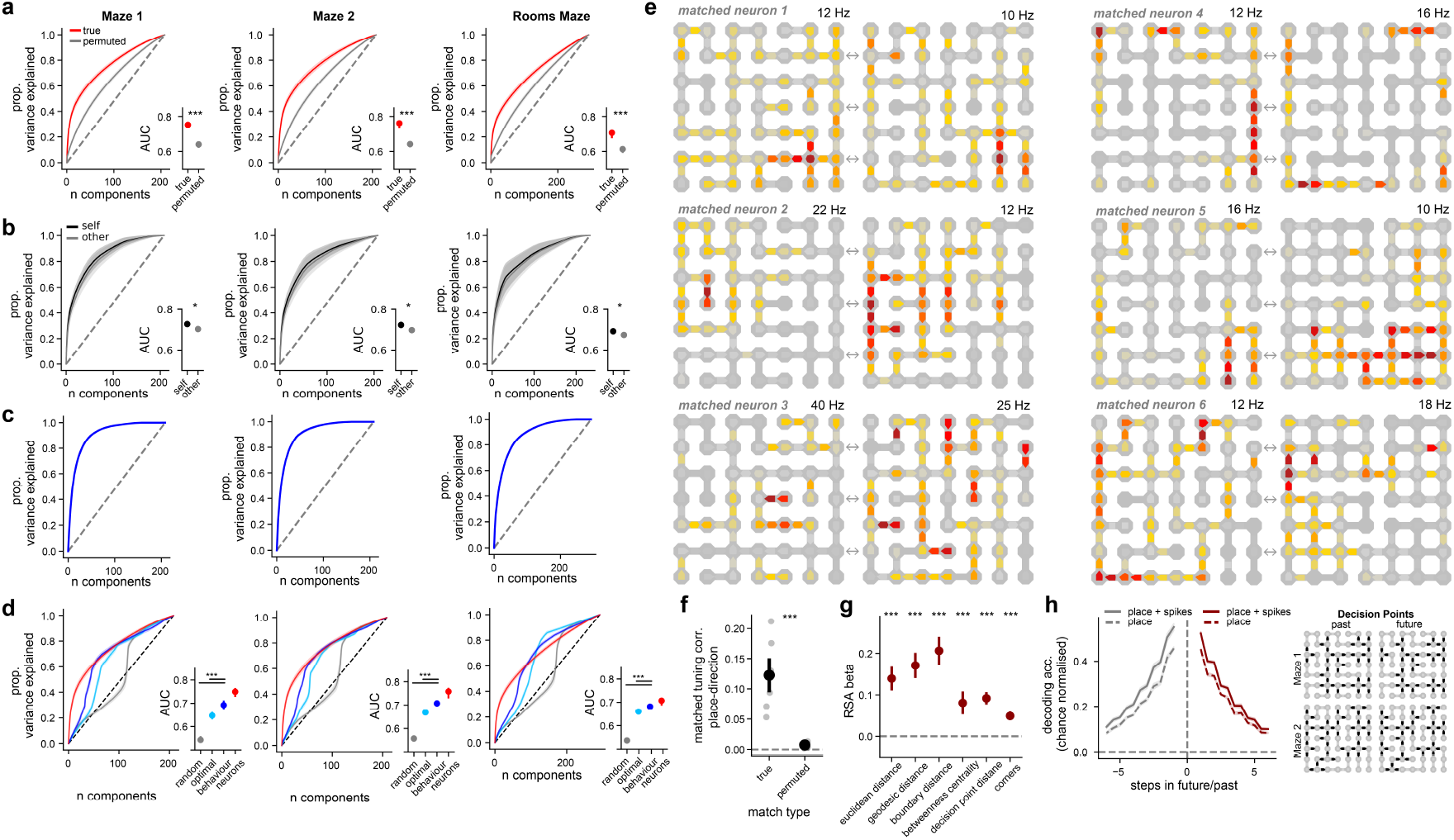
Conjunctive tuning to maze place and movement direction. **(a)** To verify that the observed place-direction tuning patterns were not artefacts of neural and behavioural autocorrelation, we compared how quickly the neural principal components (PCs) explained variance in real (red) place-direction heatmaps versus heatmaps built from neural activity circularly permuted relative to behaviour (permuted, grey). The area under each cumulative-variance-explained curve (AUC), visualised as 95% confidence intervals, was completely non-overlapping between conditions (p<.001, all mazes). **(b)** To assess whether the low-dimensional structure of place-direction tuning was shared across subjects, we quantified how well principal components (PCs) derived from the held-out subject’s own population place-direction tuning, or from all other subjects, explained variance in the held-out subject’s data. PCs from other subjects explained variance nearly as rapidly as the subject’s own, indicating strong correspondence in the low-dimensional structure across subjects. AUC (right) are formatted as in panel a (paired: t-tests: Maze 1: *T* (5) = 3.22 *p* = .023, Maze 2: *T* (5) = 4.31 *p* = .008, Rooms Maze: *T* (5) = 2.87 *p* = .037). **(c)** Cumulative variance explained curve from PCA on the behavioural data matrix comprising subjects’ navigation trajectories represented as binary place-direction vectors, reveals low-dimensional structure. **(d)** Same neuro-behavioural variance explained analysis described in Figure 4m for all mazes. Formatted as in the main-text figure. **(e)** Example neurons recorded across mazes that were tuned to: decision points (top left), corridors moving south (middle left), dead-ends flowing into the central parts of the maze (bottom left), and border-routes around the periphery of each maze moving clockwise or anti-clockwise (right). **(f)** To test if place-direction tuning remapped non-randomly across mazes, we compute the average place-direction tuning correlation (over place-direction pairs that existed on both mazes) for UnitMatch outputs (true cross-maze matches) and permuted matches (random pairs of place-direction tuned neurons recorded across mazes). The true match correlation was much higher than the permuted controls across subjects, suggesting that place-direction remapping was highly non-random. **(g)** We performed representational similarity analysis (RSA), comparing the neural RDM across maze locations to a set of candidate model RDMs derived from spatial and structural features of the maze: Euclidean distance, shortest-path distance, boundary distance, betweenness centrality, distance to decision points, and corner identity (see Methods). All features explained significant variance in neural place-direction tuning. Plotted: mean± sem RDM weights across subjects, grey markers indicate individual subject weights (one-sample *t*-tests *>* 0, BH-corrected, p < .05). **(h)** Raw decoding accuracy of decoders trained to predict subject’s past/future place-direction at decision points during navigation from subject’s current place-direction (place-dir; dashed line) or subject’s current place-direction and instantaneous neural spike vectors (place-dir. + spikes; full line). Right: “Decision points” for “past” and “future” references, visualised on Mazes 1 & 2.

**Figure S6:**
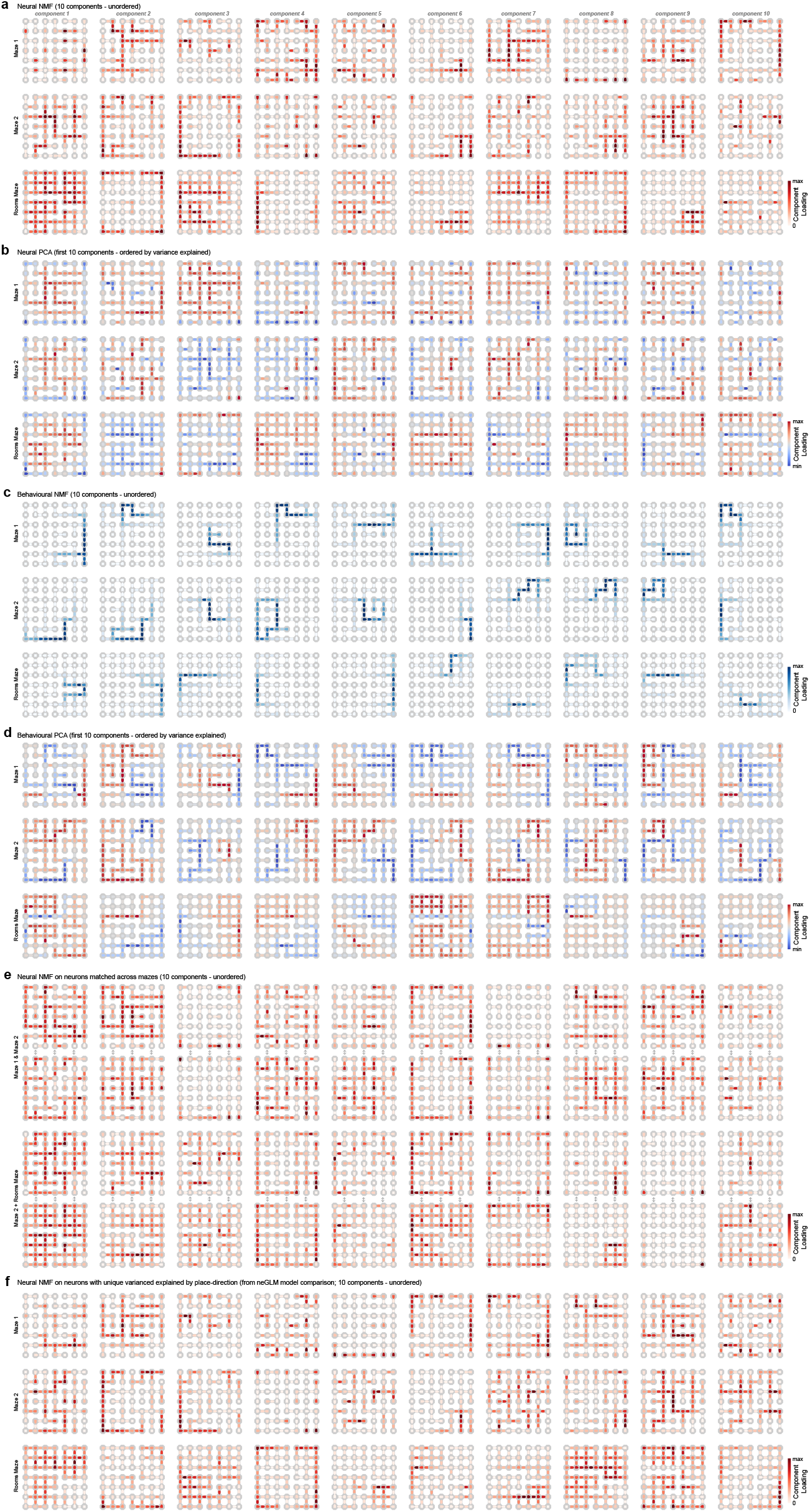
Dimensionality reduction on population place-direction tuning and navigational trajectories. **(a)** All components from a 10-component non-negative matrix factorisation (NMF) decomposition of population place-direction heatmaps (unordered). Row 1: Maze 1, row 2: Maze 2, row 3: Rooms Maze. **(b)** First 10 principal components from PCA decomposition of the same neural place-direction data matrix. Formatted similarly to a. **(c)** All components from a 10-component NMF on all navigation trials represented as place-direction sequences. **(d)** First 10 principal components from PCA decomposition of the same behavioural data matrix. **(e)** All components from a 10-component NMF decomposition of place-direction heatmaps of neurons matched across mazes. Formatted similarly to a, but each component is now represented over two heatmaps: top and bottom of each row. **(f)** Same decomposition as (a), but restricted to neurons with significant unique variance explained by place-direction (neGLM model comparison).

**Figure S7:**
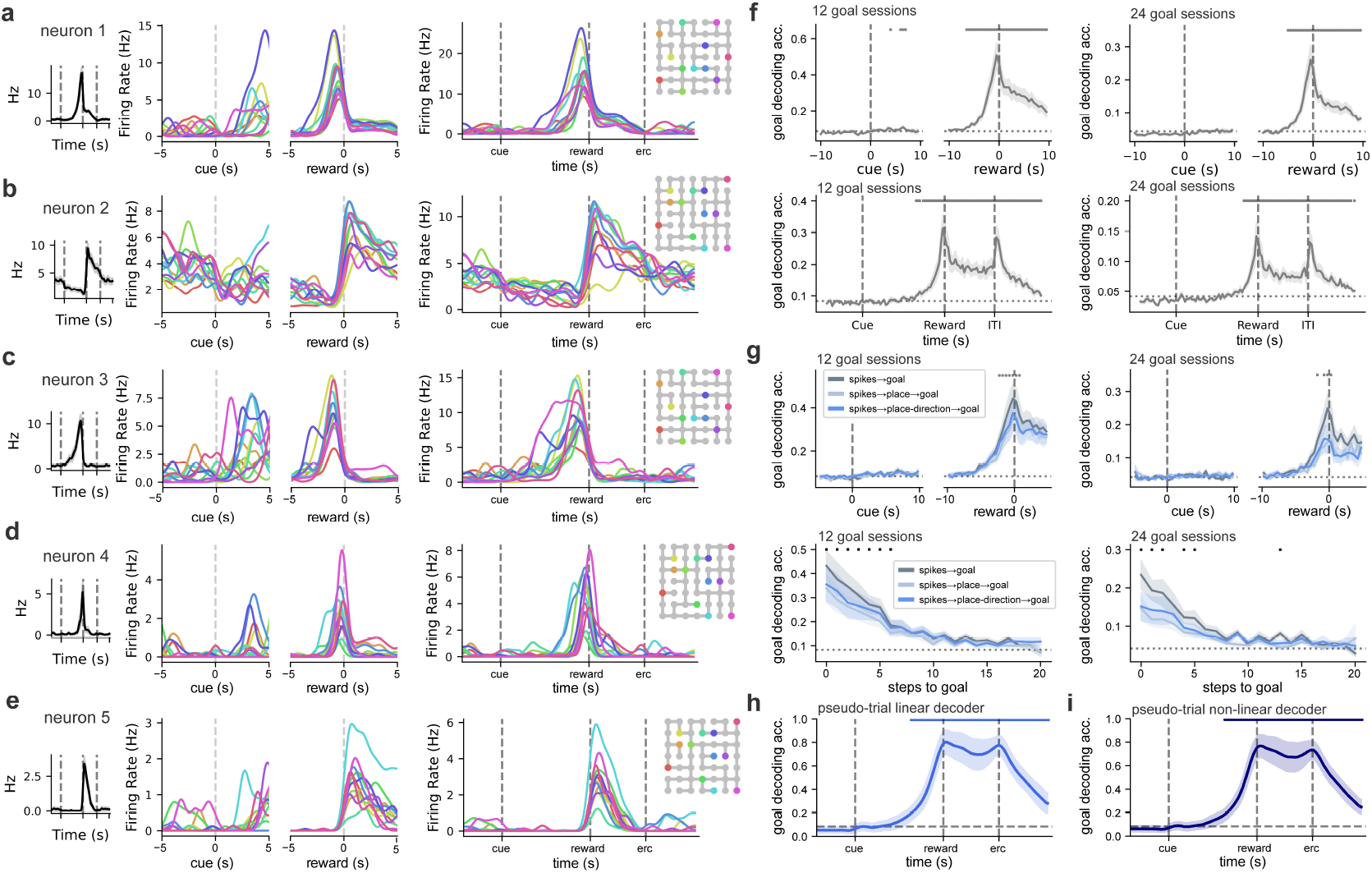
Allocentric goal decoding. **(a-e)** Event-aligned (left column) and trial-aligned (right column) firing rates of single neurons coloured by navigational goal (coloured according to maze legend in rightmost section of each panel). Goal-averaged trial-aligned response shown in leftmost section of each panel. **(f)** Event-aligned (top row) and trial-aligned (bottom row), single session goal decoding performance averaged across late 12-goal (left column) or late 24-goal (right column) sessions. **(g)** Allocentric goal decoding performance from spikes (grey), maze place decoded from spikes (light blue) and maze place-direction decoded from spikes (darker blue), aligned to cue and reward events (top row) or by steps to goal (bottom) averaged separately across 12-goal (left column) or 24-goal (right column) sessions. **(h-i)** Goal decoding from trial aligned neural activity combined across sessions with either linear (multinomial logistic regression, h) or nonlinear (multi-layer perceptron, i) decoders. Significance marked by asterisk; t-tests, BH-corrected *p <* .05. Shaded error represents SEM across subjects.

**Figure S8:**
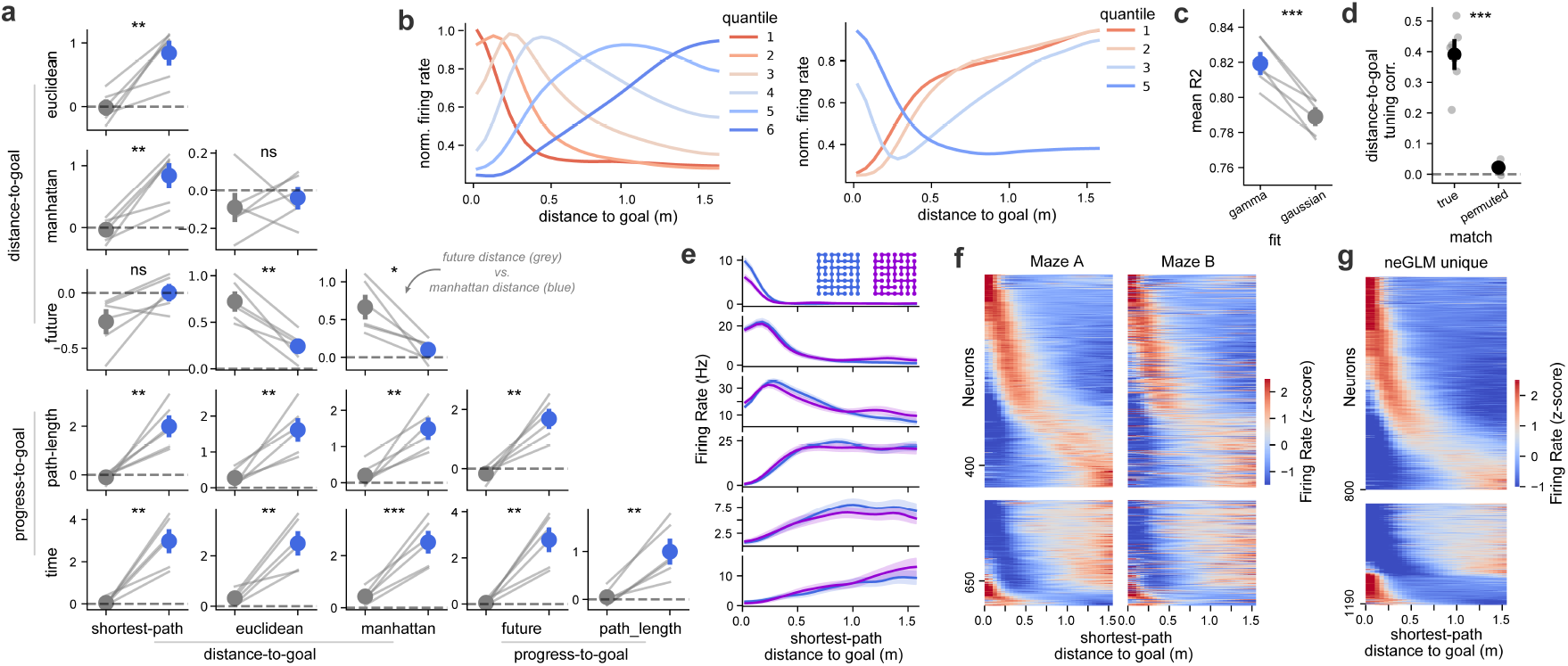
mFC represents shortest-path distance to goal. **(a)** Unique deviance explained by different distance-to-goal and progress-to-goal metrics under all pairwise comparisons. Each subplot compares distance metrics in its corresponding row and column. Coloured markers and error bars represent mean and sem across subjects. Paired t-tests for each metric comparison are shown on each subpanel (*: p<.05, **: p<.01, ***: p<.001). **(b)** Average quantiles from population distance-to-goal tuning heatmap in Figure 5f demonstrate tight tuning at close distances and broad tuning at longer distances. **(c)** To test whether population distance-tuning profiles were better described by Gaussian or gamma functions, we fit both to each distance-tuned cell on a random half of trials and evaluated goodness of fit (*R*^2^) against the tuning curves estimated from the held-out half. The gamma function best fit the data (*T* (5) = 7.322, *p <* .001) **(d)** To test if distance-to-goal tuning remapped non-randomly across mazes, we computed the average tuning-curve correlation (over place-direction pairs present on both mazes) for true cross-maze matches (UnitMatch outputs) and permuted matches (random pairs of distance-to-goal-tuned neurons recorded across mazes). The true match correlation was much higher than the permuted control across subjects, suggesting that distance-to-goal remapping was highly non-random. **(e)** Example neurons recorded on both Maze 1 (blue) and Maze 2 (purple) with similar distance-to-goal tuning on both mazes. **(f)** Coherent distance-to-goal tuning across mazes visualised as a heatmap. Distance-tuned cells were shuffled into left (Maze A) and right (Maze B) heatmaps where both heatmaps were ordered by the maximum of the gamma-fit in heatmap A (or minimum in the case of “negative” tuned cells; bottom block). **(g)** Population distance-to-goal tuning heatmap, formatted as in Figure 5f but restricted to neurons with significant unique variance explained by distance-to-goal (neGLM model comparisons), demonstrates that neurons tuned to near, intermediate, and far distances robustly encode distance to goal.

**Figure S9:**
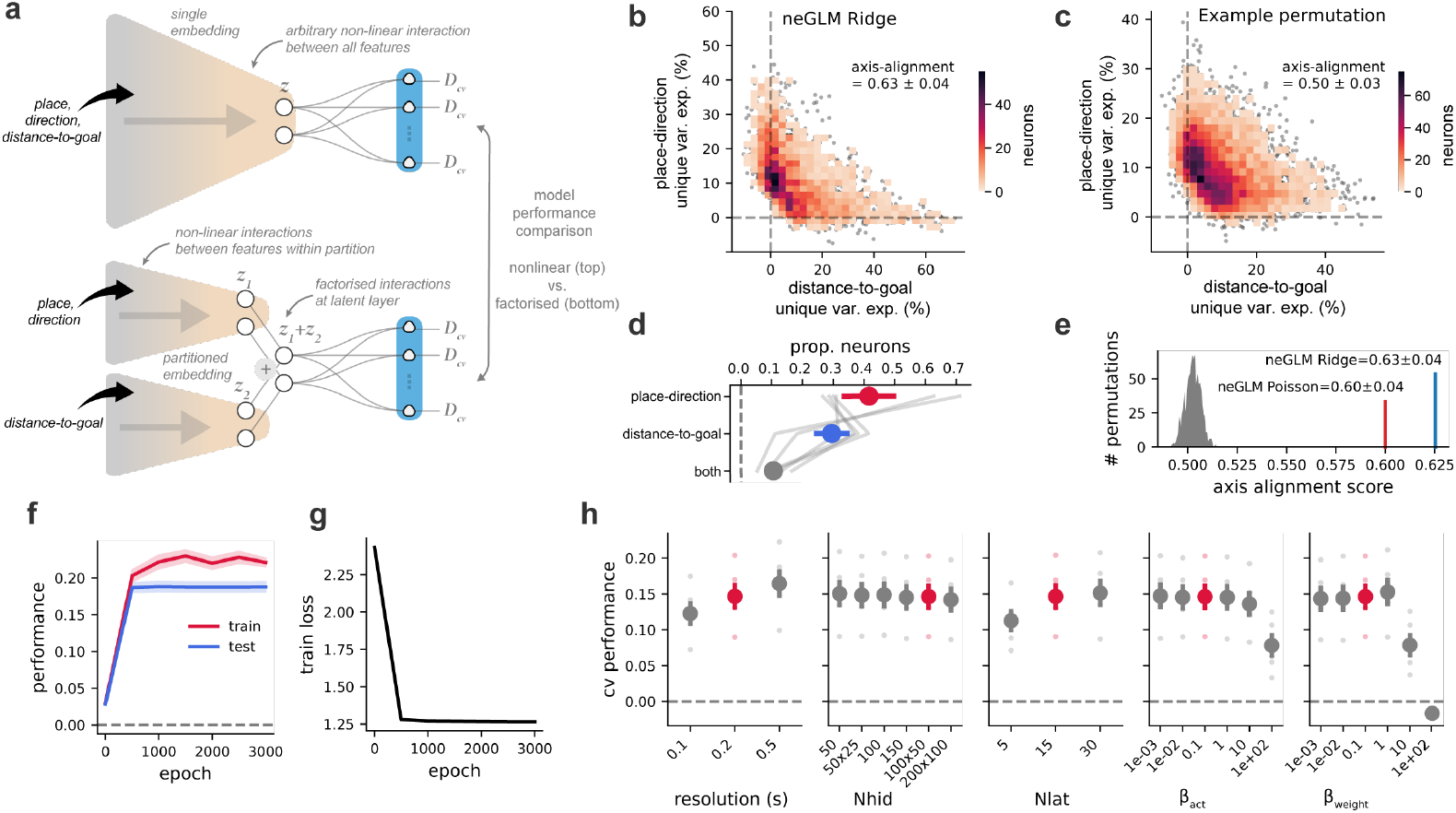
The neural embedding GLM (neGLM) **(a)** Schematic illustration of model comparisons with partitioned (facilitating factorised interactions) and non-partitioned (standard; nonlinear variable interactions) embeddings. In the non-partitioned neGLM (top), the embedding network learns arbitrary non-linear combinations of place, direction and distance-to-goal in latent units (*z*) that linearly combine to predict firing rates of held-out data. In the partitioned neGLM (bottom), the top partition learns non-linear combinations of place and direction (*z*_1_) and the bottom partition a nonlinear compression of one-hot distance-to-goal features (*z*_2_); these are then summed in the latent layer, forcing a factorised interaction between partitioned variables to predict neural activity. **(b-c)** To test whether place-direction and distance-to-goal representations were carried by largely separate sets of neurons, rather than being mixed across the population, we compared the empirical axis-alignment score – which quantifies how strongly unique variance explained clusters along the variable axes – against a null in which the held-out spikes were randomly rotated across neurons. Because rotated spikes are real-valued and incompatible with the original neGLM Poisson readout, both the null and its matched reference were computed with a Ridge (Gaussian-likelihood) regression. Unique variance from the Ridge readout (b; formatted as Figure 6g) was similar to that from the original Poisson readout (Figure 6g), clustering primarily along each variable axis (axis-alignment score: 0.63±0.04). Under the spike-rotation permutation, which enforces mixed representations across neurons, unique variance was less spread along the variable axes (example permutation in c; formatted as Figure 6g), captured by lower axis-alignment scores (0.50±0.03) across subjects. **(d)** Proportion of neurons with significant unique variance explained for place-direction, distance-to-goal or both variables (both) across subjects from neGLM model comparisons. Significance determined for each variable by a *t*-test *>* 0 of unique variance explained across folds (see Methods). **(e)** Null distribution of the subject-mean axis-alignment score across 1,000 random rotations (grey). Coloured markers indicate the observed values under the Poisson (red) and Ridge (blue) readouts on the un-rotated (empirical) data. Both fall far outside the null, indicating that place-direction and distance-to-goal information is carried by largely distinct subpopulations of neurons rather than mixed across them. **(f-g)** Embedding network training dynamics (neGLM step 1). Training loss (f) and train vs held-out-test embedding performance (g) across training epochs (sampled every 500 epochs) for the baseline neGLM model described in h. Train and test performance track closely and plateau well before the end of training, indicating convergence without overfitting. **(h)** neGLM hyperparameter sweep. Performance of the neGLM models on Maze 1 with input variables: place, direction and distance-to-goal (non-partitioned) as a function of each model hyperparameter, including: input spatial resolution, encoder hidden-layer sizes (*N*_*hid*_), latent dimensionality (*N*_*lat*_), activity regulariser (*β*_*act*_) and weight regulariser (*β*_*weight*_). Within a given hyperparameter sweep all other hyperparameters are held at their default values (red).

**Figure S10:**
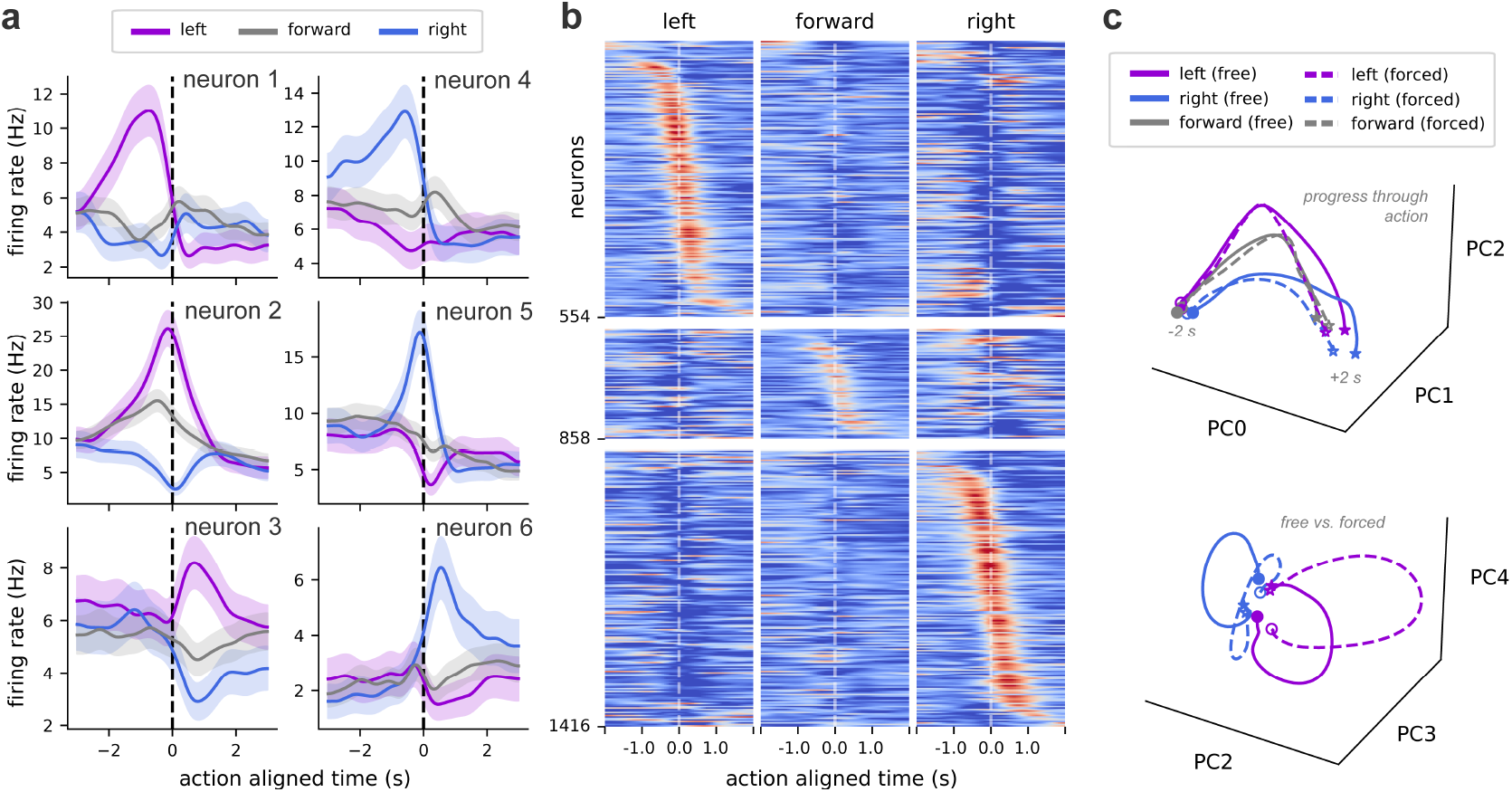
Neural tuning to egocentric actions. **(a)** mFC neurons fire selectively for different egocentric actions (‘turn left’: purple, ‘turn right’: blue, ‘go forward’: grey) relative to tower entry (0 s), where some cells increase their firing rate just before turning left (top-left panel) and some neurons fire just after turning right (bottom-right panel), for example. **(b)** Visualising this tuning across the population shows that mFC tiles progress through egocentric action. Each row is a single neuron, with columns split into blocks showing tuning to left, forward and right actions. Rows are grouped by preferred action (top: left, middle: forward, bottom: right) and ordered within each group by the time of maximum firing rate relative to the preferred action. **(c)** These population dynamics can also be visualised by projecting average population activity for each egocentric action into the low-dimensional space capturing the most variance between action-types, further stratified into ‘free’ (towers with 3 or 4 connected bridges, i.e. decision points) and ‘forced’ (1 or 2 connected bridges) actions. Top: PC1, PC2 & PC3 capture progress through different egocentric-actions. Bottom: PC4 & PC5 appeared to differentiate between free and forced egocentric actions. *Note*: Although egocentric action explains only a small amount of unique variance in the neGLM model comparisons, this effect may be systematically underestimated. Tuning to egocentric action could arise from nonlinear interactions between place, allocentric direction, and distance to goal without an explicit egocentric-action regressor. Neurons here were not selected for unique variance explained by egocentric-actions (neGLM model comparisons), but for stable action-aligned tuning curves (see Methods).

**Figure S11:**
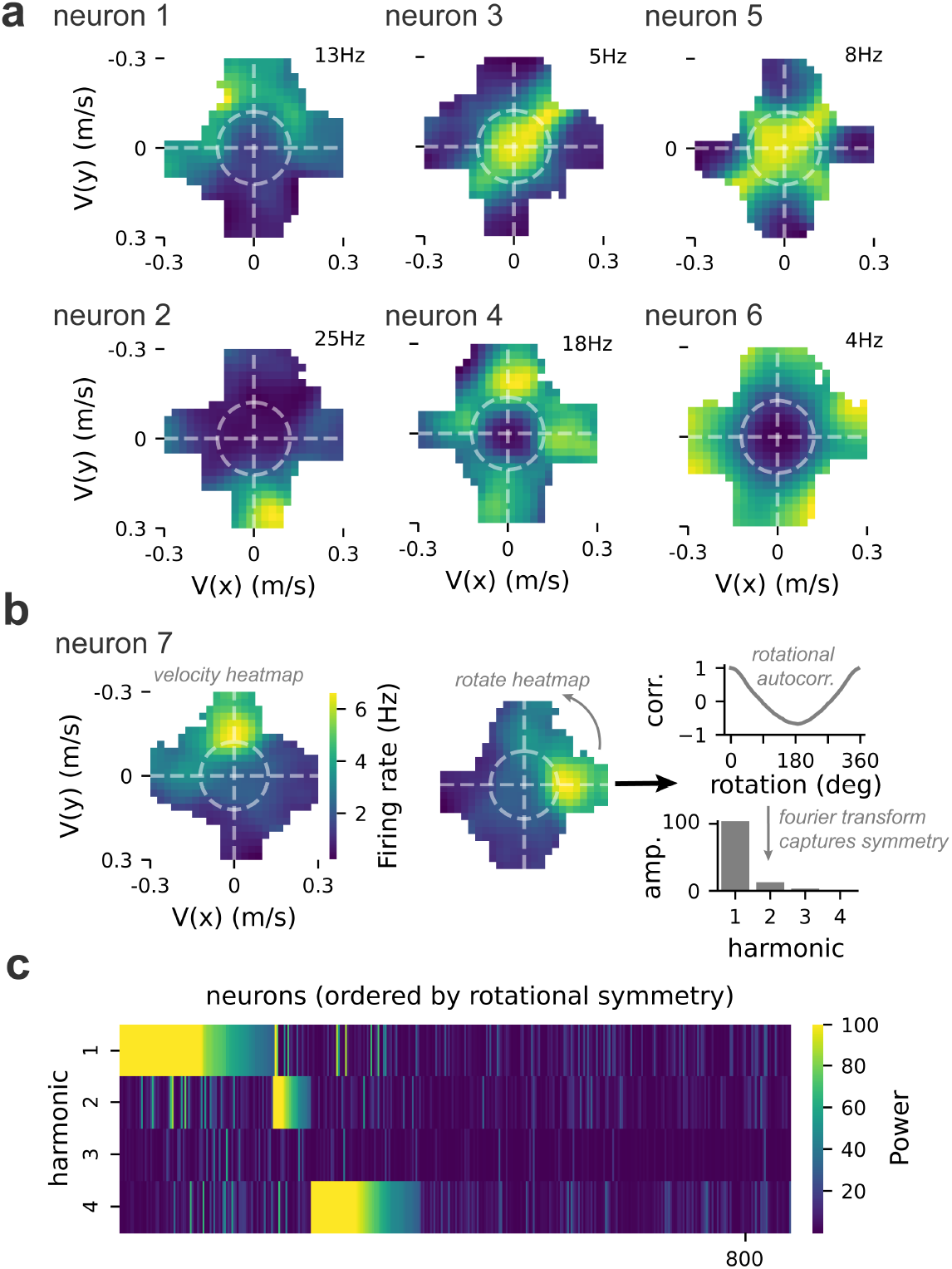
Neural tuning to movement velocity. **(a)** Individual mFC neurons showed diverse tuning to movement velocity, visualised as 2D firing-rate maps over *x* and *y* velocity. Tuning was heterogeneous across the population: some neurons fired most strongly during fast movement in a preferred direction (neurons 1–2), others showed multi-peaked tuning along on- or off-cardinal axes (neurons 3–5), and others fired with movement speed regardless of direction (neuron 6). Each neuron’s maximum firing rate is shown top right (colorbar in b). Dashed white lines, cardinal axes; dashed white circle, movement threshold (*>* 0.12 m*/*s). **(b-c)** To summarise velocity tuning across the population, we computed the rotational autocorrelogram of each neuron’s velocity heatmap and took its discrete Fourier transform; the resulting spectrum captures the rotational symmetry of the tuning field, with the dominant harmonic giving its *n*-fold order (pipeline in b). Plotted across neurons (c), this revealed three groups: a large population with low power across all harmonics, consistent with pure speed modulation; a population dominated by the 1st harmonic, corresponding to single-peaked velocity fields (e.g., neurons 1 & 2); and a population dominated by the 4th harmonic, corresponding to on/off cardinal axis tuning (e.g., neurons 4 & 5). *Note*: The above population analysis includes cells selected for stable velocity tuning curves (see Methods), not for unique variance explained by velocity in neGLM model comparisons. As such, some patterns in velocity tuning may be derivative of tuning to other represented variables (e.g., place-direction or distance-to-goal).

**Figure S12:**
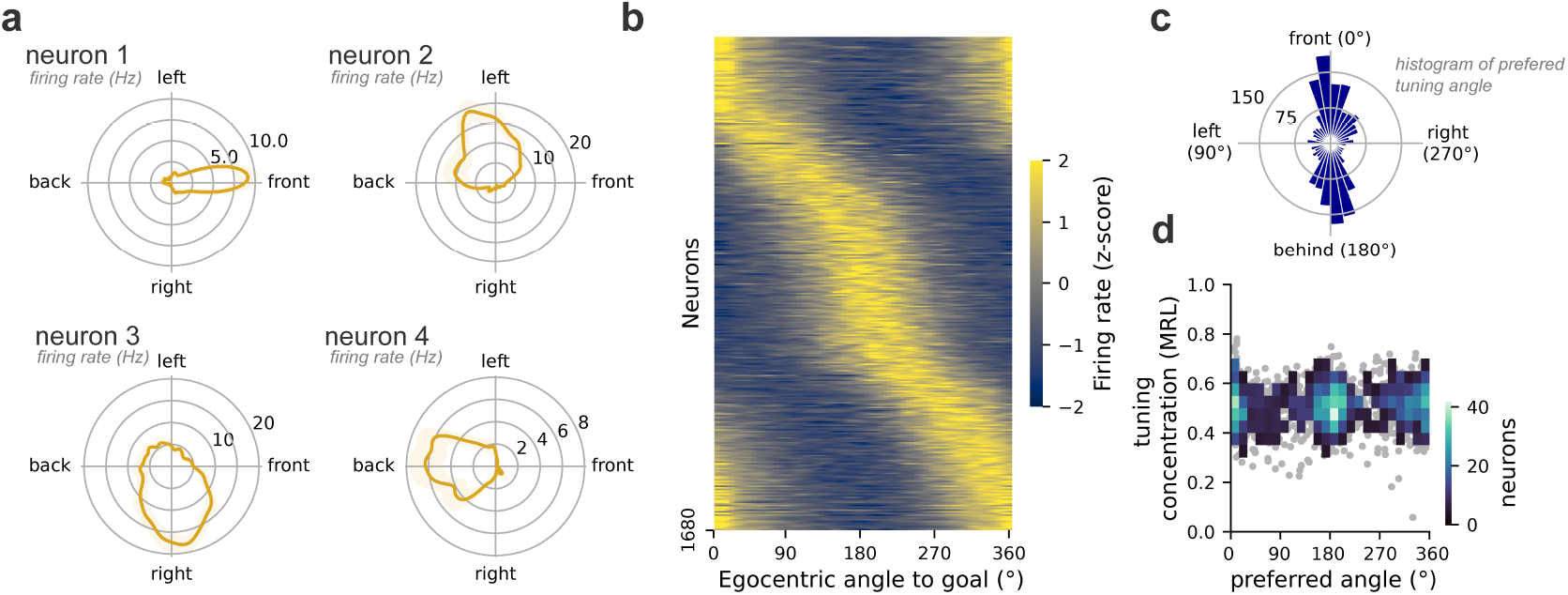
Neural tuning to egocentric-angle-to-goal. **(a)** Polar tuning curves for four example single units, each preferring the goal in a different egocentric direction (front, left, right, back). Radial axis: mean firing rate (Hz); shaded area represents SEM across trials. 0° = goal directly ahead of the animal’s current heading; angle increases counter-clockwise. **(b)** Population tuning-curve heatmap; neurons sorted by preferred angle (cross-validated von Mises fits). Two dense bands are visible, corresponding to goal-ahead (0°) and goal-behind (180°) sub-populations. **(c)** Circular histogram of preferred angles for the same neurons. **(d)** Joint distribution of preferred angle (x-axis) and tuning concentration (mean resultant length, MRL; y-axis; 0 = uniform, 1 = sharply tuned). Grey dots: individual neurons (shown so outlier bins remain visible); coloured bins: 2D histogram with sparse bins masked. *Note*: Neurons here were not selected for unique variance explained by egocentric-angle-to-goal (neGLM model comparisons), but for stable angle-to-goal tuning curves (see Methods). As such, some patterns in the population tuning above may be derivative of correlated variables (e.g., distance-to-goal).

**Figure S13:**
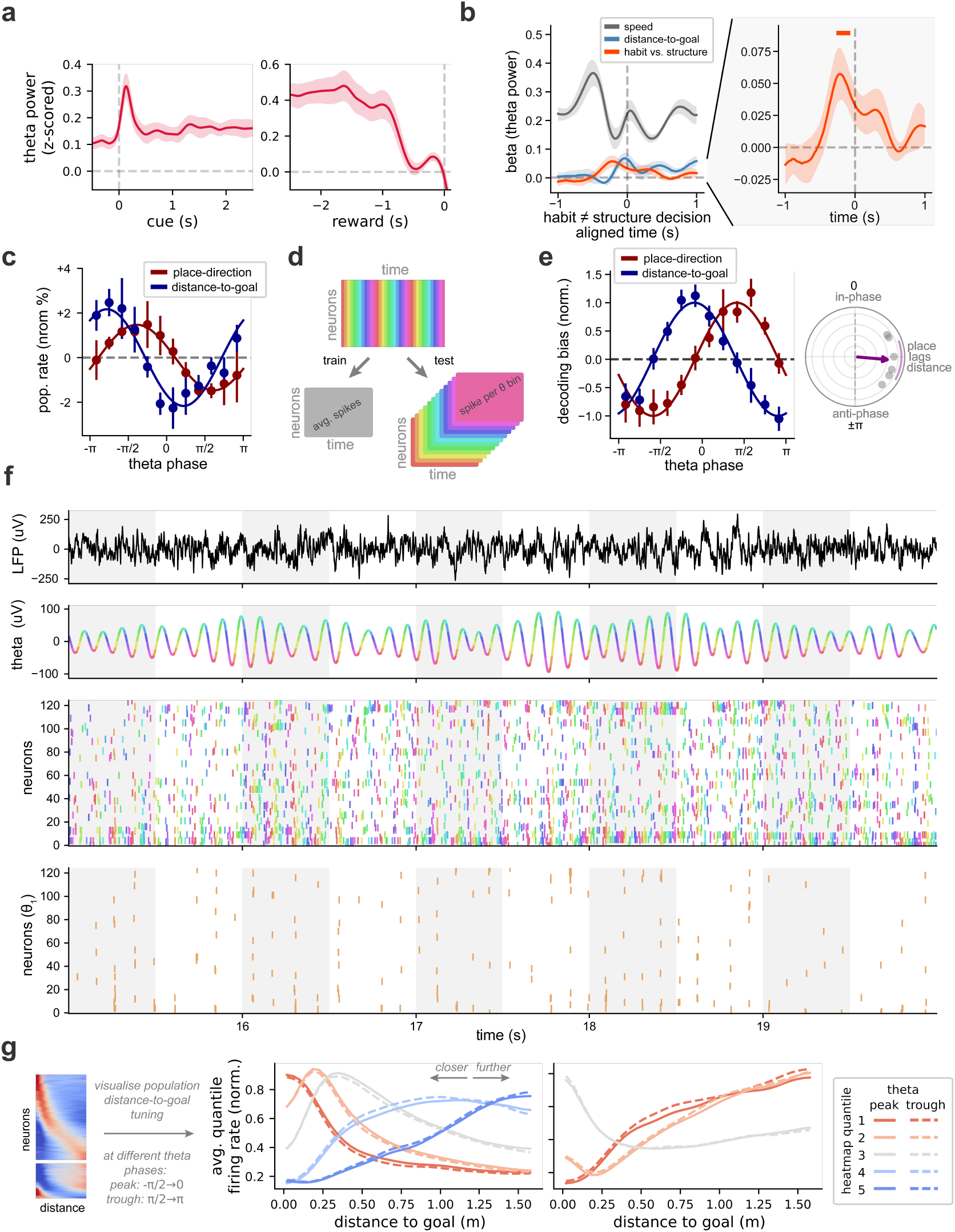
Theta-modulated representations in mFC. **(a)** Theta power (z-score normalised) aligned to trial events: cue (left) and reward (right). **(b)** Structure ≠ habit decision aligned linear regression weights for speed, distance-to-goal, and a choice (habit vs. structure) to explain theta-power. Weights fit separately for every timepoint. Left: enlarged time-course of choice regressor, showing significantly stronger theta power before structure-based decisions. Solid orange bars indicate timepoints where cross subject weights were *>* 0 (*t*-tests > 0, without multiple comparisons correction). **(c)** Average modulation of population firing rates for neurons with significant variance explained by either place-direction or distance-to-goal. Formatted as in Figure 7b-bottom. **(d)** Schematic illustration of theta-stratified logistic regression decoding approach. Spikes are stratified by theta phase (12 bins) across the session, yielding 12 independent neuron-by-timepoint matrices (colour = theta-phase bin). Critically, each matrix can be downsampled to a lower resolution (greatly improving decoding performance) while preserving phase-specific information (see f). **(e)** Train/test sample matched distance-to-goal and place-direction theta-stratified decoding biases curves (left), with cross subject phase-offset (right). Formatted as in Figure 7e-i. **(f)** Further illustration of the spike-by-theta phase stratification for theta-modulated decoding and tuning curve analyses. Top: raw LFP. Second row: the same LFP band-pass filtered in the theta band (7–11 Hz); each segment of the filtered trace coloured by one of twelve theta-phase bins. Third row: spike raster across isolated single and multi units; each spike is coloured by the theta-phase bin in which it occurred. Bottom row: the same raster showing only spikes from a single phase bin. Faint vertical grey bands indicate 0.5 s windows over which spikes per theta-phase can be summed when downsampling the data. **(g)** To independently verify theta-modulation of the distance-to-goal representation (shown initially using a decoding approach in Figure 7d), we compared distance-to-goal tuning curves across theta-phases. For each distance-tuned cell (heatmap, left), we derived a distance-to-goal tuning curve from spikes in each of 12 theta-phase bins, plus a reference curve from activity averaged over all phases. Stacking these curves across neurons gave 12 phase-specific population distance-tuning matrices (+1 reference). We then found the x-shift (in distance-to-goal) that best aligned each phase-specific matrix to the reference, by minimising mean squared error. This revealed systematic modulation of the distance representation over theta-phase (Figure 7e). For further visualisation, we computed population tuning quantiles from the distance-tuning matrices (heatmaps) at the peak and trough of theta phase. Formatted as in Figure S8b, with peak-spike quantiles as full lines and trough-spike quantiles as dashed lines.

## Footnotes

1 ”on one occasion he was discovered on the local golf course having originally stepped outside the therapy room to fetch some coffee” — Shallice & Burgess describing a patient with bilateral damage to the prefrontal cortex [1]

